# Analysis of gene expression profiles to elucidate racial differences in African American and White patients with Triple-negative breast cancer

**DOI:** 10.1101/2024.05.29.596478

**Authors:** Hailey McAndrew, Jessica Rigler, Suneetha Yeguvapalli, Kumaraswamy Naidu Chitrala

## Abstract

Triple-negative breast cancer (TNBC) is the second most diagnosed subtype of breast cancer. It is known to be the most aggressive one that lacks known targetable receptors. One of the concerns in TNBC is the disparities in its prevalence and tumor pathogenesis among women with non-Hispanic African American backgrounds. Despite extensive research, the genetic underpinnings that lead to these disparities remain elusive. The current study aims to provide initiative for further clinical research in the development of targeted therapy for TNBC. Gene expression profiles from African American (AA) and European American (EA) patients with TNBC were collected from Gene Expression Omnibus and performed differential gene expression (DEG)analysis. Candidate genes for a significant correlation between expression and survival rates for breast invasive carcinoma were analyzed using UALCAN. The DAVID annotation tool, Enrichr web server, KEGG database, and Gene Ontology (GO) database were used for functional enrichment analysis of target genes. The Network Analyst server was used to identify ligands with strong affinities, SeamDock server for molecular docking between the biomarkers/associated ligands and examined protein-protein interactions (PPI) from the STRING server. Data from public breast cancer cohorts was utilized to identify expression patterns associated with poor survival outcomes of AA patients with TNBC. Our results showed three genes of interest (*CCT3*, *LSM2*, and *MRPS16*) and potential ligands for molecular docking. Molecular docking was performed for the ICG001 ligand to *CCT3* (binding affinities of -9.3 kcal/mol and -8.9 kcal/mol) and other interacting proteins (*CDC20* and *PPP2CA*) with high degrees of connectivity. The results determined molecular docking of ICG001 to the *CDC20* protein resulted in the highest binding affinity. Our results demonstrated that *CCT3* and its interacting partners could serve as potential biomarkers due to their association with the survival outcome of AA patients with TNBC and ICG 001 could be the therapeutic lead for these biomarkers.

## Introduction

Breast cancer is the leading cause of cancer death in women. Female breast cancer accounts for 11.7% of all cancer cases, surpassing lung cancer at 11.4% [1]. According to the GLOBOCAN statistical review, in 2020 there were approximately 2,261,419 new breast cancer cases worldwide. The global burden of incidence and mortality rates due to breast cancer continues to increase at an alarming rate [1].

Breast cancer is a heterogeneous disease that is known to manifest a wide range of diverse clinical presentations. The current breast cancer diagnostic procedure involves a series of medical evaluations, including screening, tissue biopsy, and laboratory testing [2]. Additional laboratory testing is performed following a diagnosis to assess expression levels of the major biomarkers associated with breast cancer. Breast cancer is classified into several molecular subtypes based on the expression of receptors: estrogen receptor (ER), progesterone receptor (PR), and human epidermal growth factor receptor 2 (HER2) [3]. Based on the presence or absence of these receptors, breast cancer tumors are classified into one of the major breast cancer subtypes: Luminal A, Luminal B, HER2-enriched, or triple-negative breast cancer [4]. Current standard treatment options available include surgery, radiation therapy, and chemotherapy, but triple-negative tumors are highly unresponsive to standard therapy rendering many conventional treatment options ineffective.

Triple-negative breast cancer (TNBC) is the second most common breast cancer subtype, representing approximately 15-20% of all breast cancer cases [5]. TNBC lacks expression of all three target receptors [6], making it challenging to accurately diagnose TNBC and limiting available treatment options. The dynamic phenotypes of TNBC often lead to diagnostic delays that compromise the ability to detect TNBC at an earlier stage when the disease is most treatable. Some clinical characteristics frequently observed in TNBC include rapid metastatic progression, greater resistance to standard therapy options, and significantly higher mortality rates than other forms of breast cancer [7]. The relative survival rate of TNBC is nearly 20% less than that of the other subtypes and declines drastically as the cancer progresses [8]. Because TNBC has no known therapeutic targets, early detection is a critical factor in successfully treating TNBC and maximizing overall survival rates [9].

A major concern of TNBC is the distinct clinical differences observed in premenopausal African American patients compared to other ethnic groups [10]. While TNBC affects women regardless of ancestral background [11], the prevalence of TNBC is notably higher and the survival probability is significantly lower in African American patients [12]. According to some studies, African American women are nearly three times more likely to be diagnosed with TNBC than European American women [12, 13]. Additionally, African American women are more commonly diagnosed with TNBC at a younger age and at a later stage, when the cancer has already metastasized [12, 14, 15]. Current evidence suggests that the burden of TNBC in African American women is heavily influenced by a combination of biological, genetic, environmental, and socioeconomical factors [15]. Understanding the complex interactions between these factors at the molecular level could provide insight about the underlying mechanisms driving the existence of ethnic disparities in TNBC.

## Results

### Data characteristics

The GSE37751 data series comprised of microarray data from 108 primary breast tissue samples, including 14 TNBC samples. The series matrix included transcriptomic measurements for 33,298 gene probes platform that were transformed into official gene symbols. The GSE142731 dataset contained data from 42 Caucasian and African American patients with TNBC, of which 35 samples were able to be analyzed by GEO2R: 19 samples from African American TNBC patients and 16 from Caucasian TNBC patients.

### Identification of DEGs

The analysis identified 1,830 significant DEGs (*P* value < 0.05) between AA and EA patients with TNBC from dataset GSE37751 (GEO accession), of which 373 genes were downregulated and 1,457 genes were upregulated. Analysis of dataset GSE142731 identified a total of 479 DEGs, including 192 downregulated genes and 286 upregulated genes. These two datasets are displayed in Supplementary Table 2. The differential expression analysis results between African American (AA) and European American (EA) patients with TNBC for the GSE37751 dataset were represented graphically in Figure 1 including the distribution of significantly upregulated and downregulated genes. The top 20 dysregulated genes in GSE37751, as determined by absolute FC, are shown in Table 1, including genes *MAPKAP1*, *SAA2*, *ZNF800*, *SCGB2A2*, *SUPT7L*, *MEGF6*, *NPAP1*, *KMT5A*, *STEAP4*, and *ORC5*. Table 2 lists the top 20 dysregulated genes in GSE142731, including *CD207*, *MIR9-1HG*, *EPHA7*, *SPTSSB*, *IGFL2*, *ATP2B2*, *ALDH3B2*, *MIR646HG*, *WFDC1*, and *NES*.

**Fig 1.**
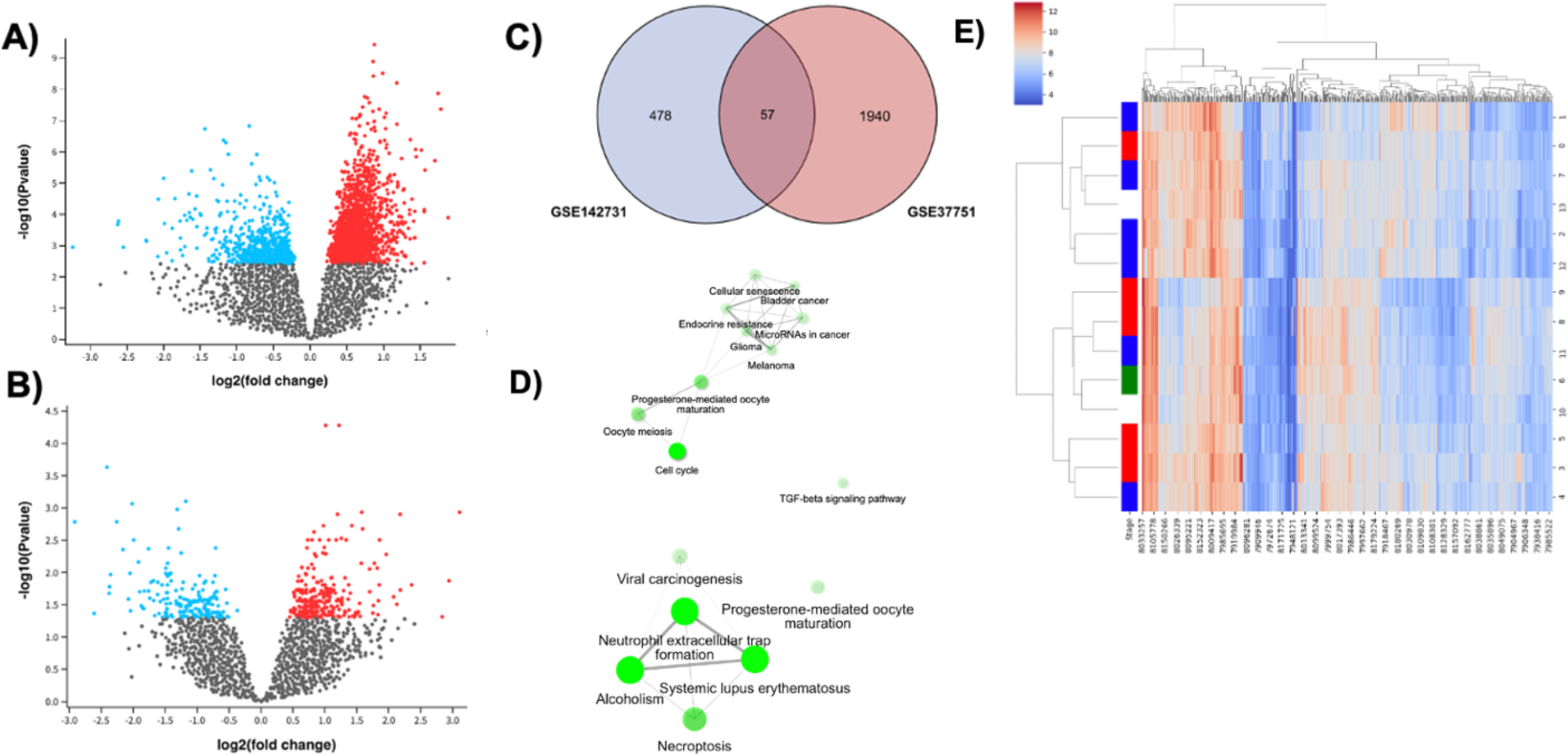
Differential expression analysis between African American (AA) and European American (EA) patients with TNBC. **(A)** Volcano plot and **(B)** Meandiff plot showing the distribution of significantly upregulated and downregulated genes between AA versus EA TNBC samples. The blue dots represent downregulated genes, and the red dots represent the upregulated genes. The black dots represent not significant genes. **(C)** Gene overlap between the two datasets. **(D)** Pathway enrichment network based on functional annotations of the identified hub genes, where the enrichment of each pathway is reflected in the opaqueness of the green nodes. **(E)** Heat map showing the expression variance of key nodes in the co-expression network in breast invasive carcinoma cells and normal cells. Gene expression profiles are displayed according to the color range shown in the key, where red represents higher expression and blue represents lower expression.

**Table 1.** Top 20 dysregulated DEGs between African American and European American patients with TNBC from dataset GSE37751. For each gene, the Entrez ID, log2 fold change (log2FC), p-value (pval), adjusted p-value (padj), and gene description are provided.

| Entrez ID | Gene | log2FC | pval | padj | Description |
| --- | --- | --- | --- | --- | --- |
| 79109 | *MAPKAP1* | -3.27 | 1.13E-03 | 2.60E-02 | mitogen-activated protein kinase associated protein 1 |
| 6289 | *SAA2* | -2.63 | 2.10E-04 | 1.05E-02 | serum amyloid A2 |
| 168850 | *ZNF800* | -2.61 | 1.64E-04 | 9.12E-03 | zinc finger protein 800 |
| 4250 | *SCGB2A2* | -2.55 | 1.13E-03 | 2.58E-02 | secretoglobin family 2A member 2 |
| 9913 | *SUPT7L* | -2.26 | 6.76E-04 | 1.97E-02 | SPT7-like STAGA complex gamma subunit |
| 1953 | *MEGF6* | -2.23 | 7.18E-04 | 2.02E-02 | multiple EGF like domains 6 |
| 23742 | *NPAP1* | -2.08 | 2.24E-03 | 3.75E-02 | nuclear pore associated protein 1 |
| 387893 | *KMT5A* | -2.07 | 3.25E-05 | 3.70E-03 | lysine methyltransferase 5A |
| 79689 | *STEAP4* | -2.00 | 7.07E-06 | 1.65E-03 | STEAP4 metalloredutase |
| 5001 | *ORC5* | -1.99 | 1.98E-04 | 1.01E-02 | origin recognition complex subunit 5 |
| 6523 | *SLC5A1* | -1.96 | 1.68E-03 | 3.19E-02 | solute carrier family 5 member 1 |
| 22861 | *NLRP1* | -1.89 | 3.06E-03 | 4.43E-02 | NLR family pyrin domain containing 1 |
| 8618 | *CADPS* | -1.88 | 2.79E-04 | 1.24E-02 | calcium dependent secretion activator |
| 4239 | *MFAP4* | -1.84 | 3.11E-05 | 3.64E-03 | microfibrillar associated protein 4 |
| 55635 | *DEPDC1* | 1.78 | 4.27E-08 | 1.19E-04 | DEP domain containing 1 |
| 124220 | *ZG16B* | -1.77 | 1.63E-04 | 9.08E-03 | zymogen granule protein 16B |
| 4969 | *OGN* | -1.77 | 5.74E-04 | 1.81E-02 | osteoglycin |
| 222235 | *FBXL13* | 1.75 | 1.34E-08 | 7.42E-05 | F-box and leucine rich repeat protein 13 |
| 10112 | *KIF20A* | -1.74 | 2.07E-03 | 3.58E-02 | kinesin family member 20A |
| 7260 | *TSSC1* | -1.71 | 1.07E-04 | 7.33E-03 | tumor suppressing subtransferable candidate 1 |

**Table 2.**
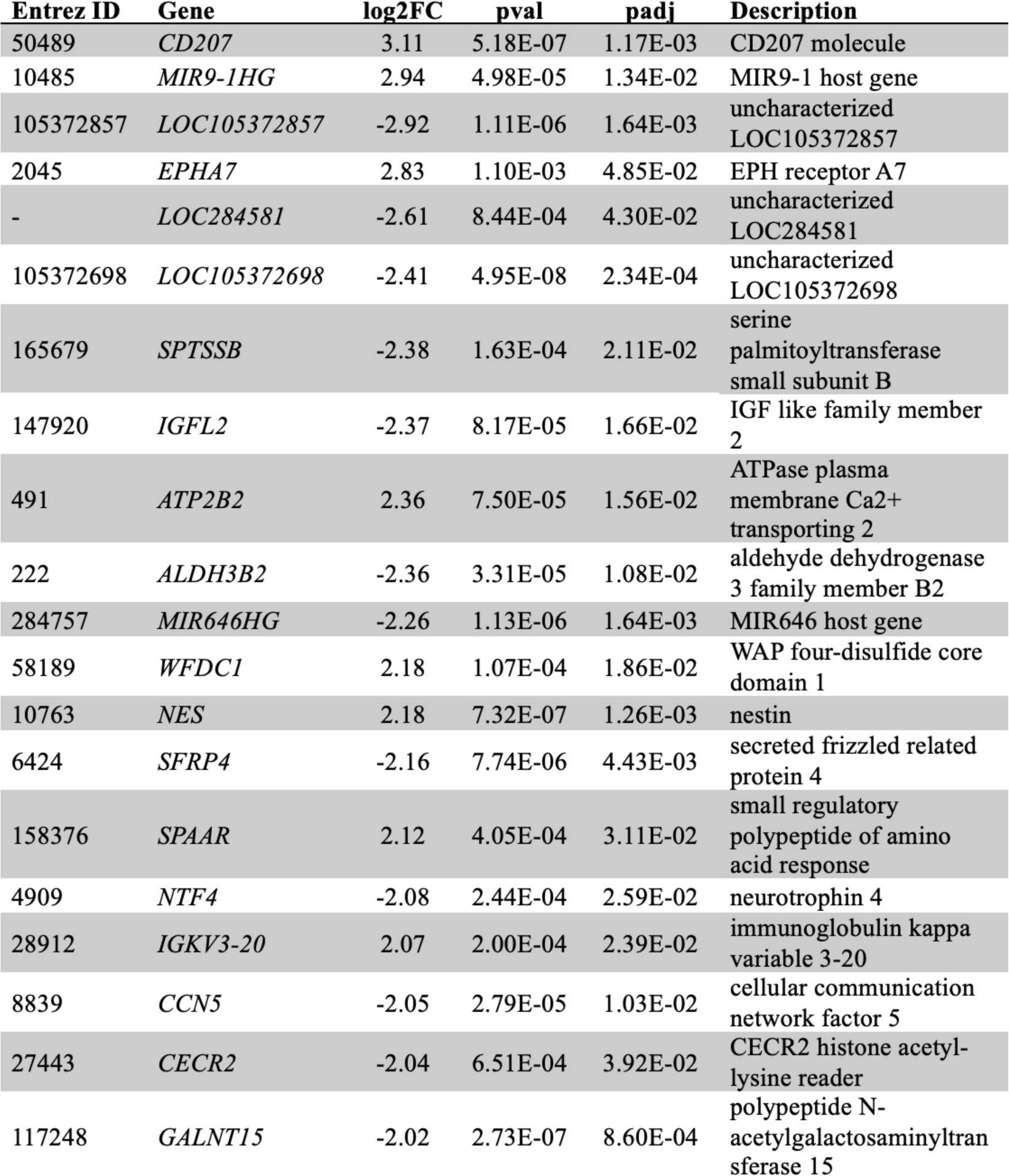
Top 20 dysregulated DEGs between African American and European American patients with TNBC from dataset GSE142731. For each gene, the Entrez ID, log2 fold change (log2FC), p-value (pval), adjusted p-value (padj), and gene description are provided.

### Identification of Potential Biomarkers

To gain further insight into the epigenetic mechanisms that contribute to race-specific phenotypes of TNBC,a systematic analysis of the DEGs from both datasets was performed using Metascape. The analysis report, depicted in Figure 2C, highlighted overlapping functional categories and pathway representation among the gene lists and mapped a protein-protein interaction (PPI) network. Three key modules were extracted from the PPI network by Minimum Common Oncology Data Elements (MCODE), consisting of genes that are upregulated in both datasets and is depicted in Figure 2D. According to the report, the modules were involved in biological processes mitotic sister chromatid segregation, translation, and U2-type precatalytic spliceosome, respectively. The top enrichment results for each module are shown in Table 5.

**Fig 2.**
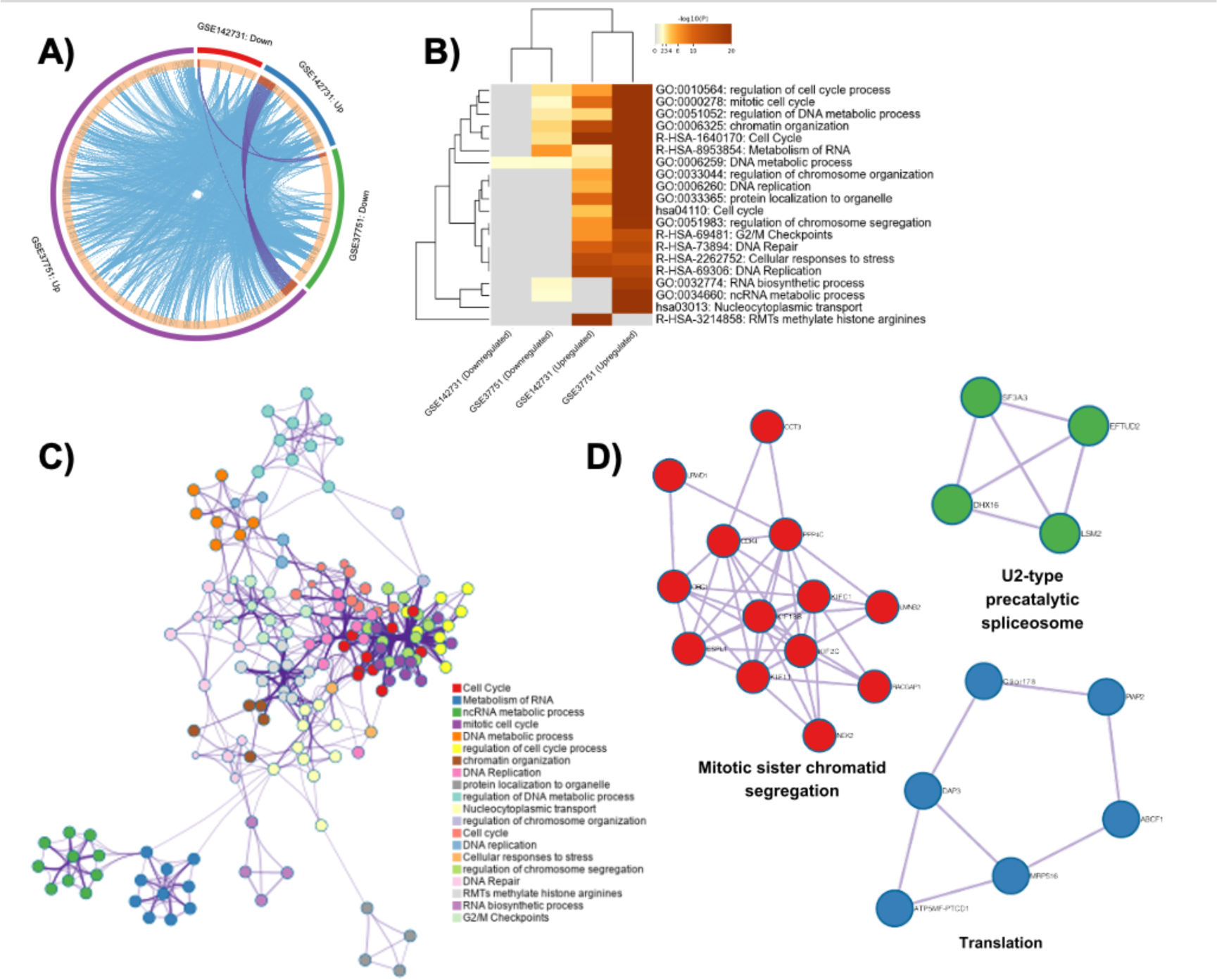
Kyoto Encyclopedia of Genes and Genomes (KEGG) pathway enrichment of differentially expressed genes (DEGs) between AA and EA patients with TNBC. **(A)** KEGG pathway enrichment plot of the upregulated and downregulated DEGs. **(B)** KEGG pathway enrichment chart of upregulated and downregulated DEGs with corresponding *P* values. **(C)** Highly connective sub-network of relevant node cluster from STRING protein-protein interaction (PPI) network of DEGs. The module consists of 17 hub genes with a high degree of connectivity (degree > 50): *CDK1, CCNB1, BRCA1, CHEK1, MCM7, MAD2L1, BUB1, AURKA, CDC20, KIF11, BUB1B, KIF23, HSPA4, ASPM, KIF20A, RACGAP1, HMMR*. **(D)** Bubble plot showing KEGG pathway enrichment of the upregulated DEGs with three modules extracted from the PPI network by Minimum Common Oncology Data Elements (MCODE).

Survival analysis of the cross DEGs from the MCODE modules against breast invasive carcinoma was performed using the UALCAN portal. The analysis found that several genes were associated with breast cancer survival, but three genes (*CCT3*, *LSM2*, and *MRPS16*) showed correlation with shorter survival times in AA patients as shown in Table 5. The survival plots for the corresponding genes are depicted in Figure 4.

**Fig 3.**
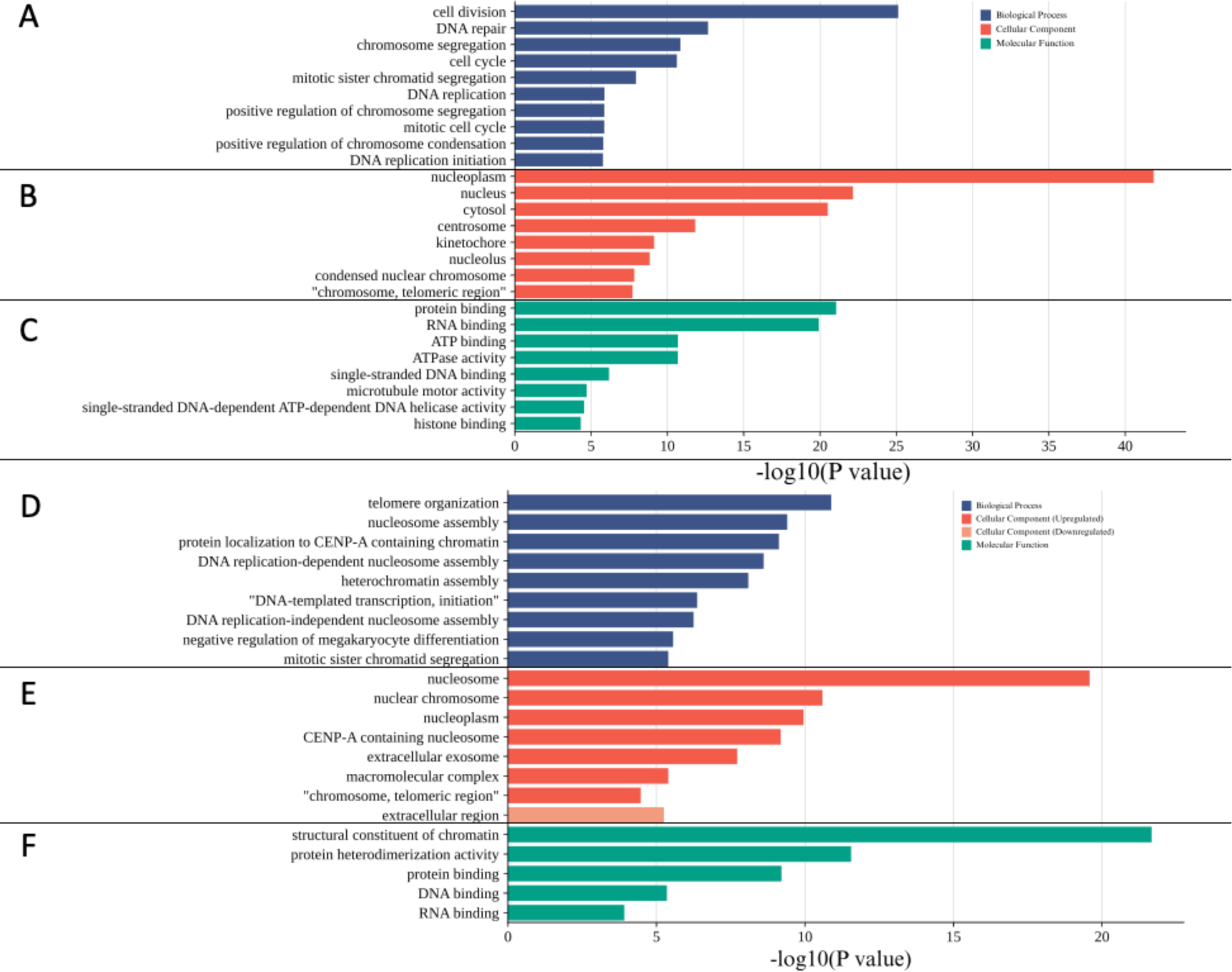
Gene ontology (GO) annotations for the upregulated and downregulated DEGs between AA and EA patients with TNBC. The bar plots show the GO terms with significant enrichment of upregulated DEGs and *P* value in **(A)** biological process, **(B)** molecular function, and **(C)** cellular component categories. The final three bar plots show the GO terms with significant enrichment of downregulated DEGs and *P* value in **(D)** biological process, **(E)** cellular component categories, and **(F)** molecular function.

**Fig 4.**
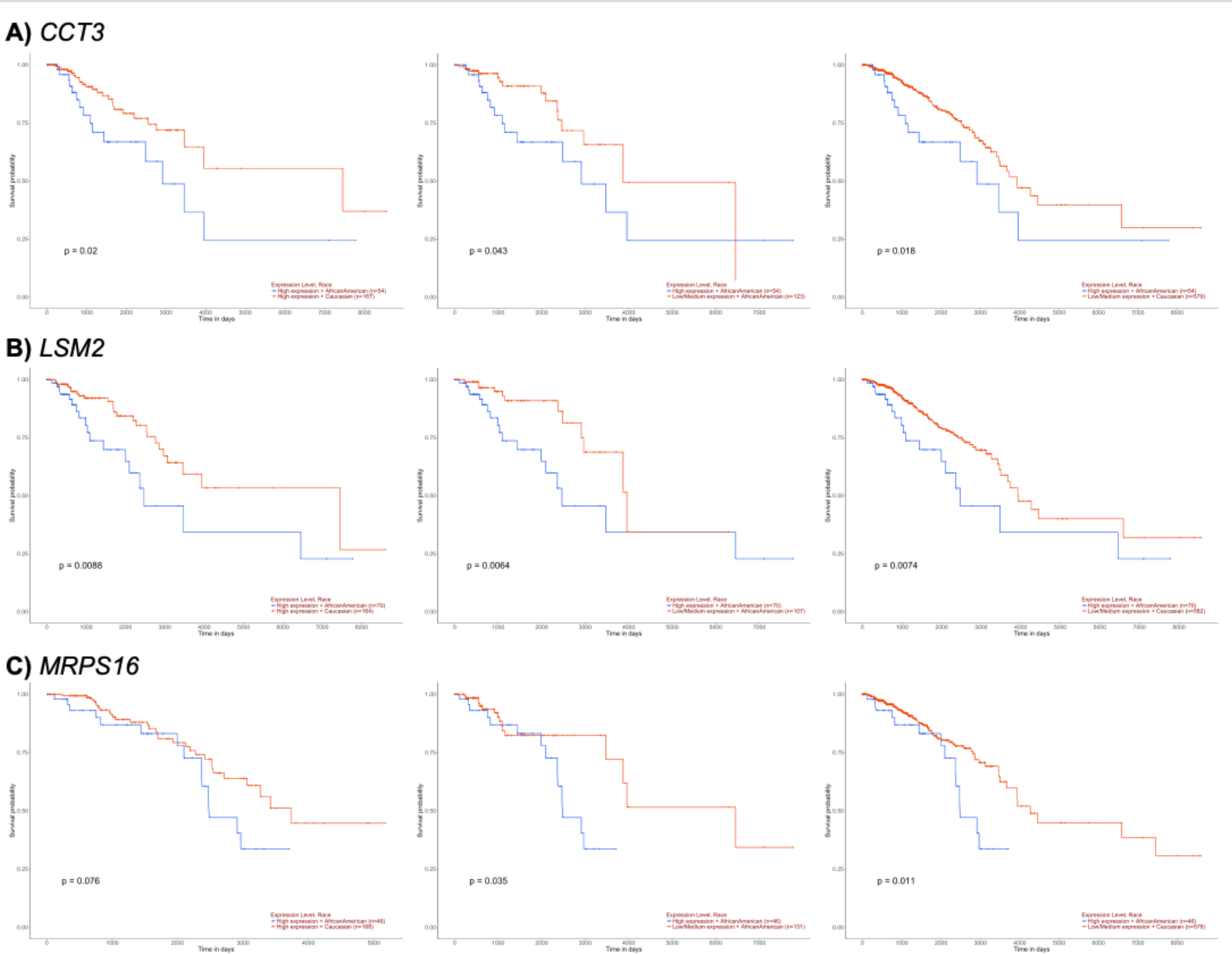
Expression profile and survival analysis of select genes in breast cancer patients. Survival plots showing the effect of **(A)** *CCT3*, **(B)** *LSM2*, and **(C)** *MRPS16* expression on patient survival for breast cancer based on patient race. The y-axis and x-axis show the survival probability (decimal) over time (in days), respectively. The first graph for each gene compares the survival probability of African American patients with high expression (red line) versus high expression with Caucasian (blue line). The second graph for each gene compares high expression with African American (red line) versus low/medium expression with African American (blue line). The third graph for each gene shows high expression with African American (red line) versus low/medium expression with Caucasian (blue line).

### Target identification and Molecular Docking

To explore potential binding targets for the genes of interest (*CCT3, LSM2, and MRPS16*), meta-analysis was performed using the Network Analyst server and are visualized in Figure 5B. The resulting ligands of interest (Atrazine and ICG 001) listed in Table 6 were prepared in PDB format using PyMol. Atrazine and ICG 001 are shared between the *CCT3* and *MRPS16* nodes as shown in Figure 5A. The SeamDock server was used to perform molecular docking. The prepared ligand file was uploaded to SeamDock and the PDB ID was entered for the receptor. The PDB ID for the *CCT3* protein was 7TTT. The molecular docking results of the *CCT3* protein with Atrazine resulted in 2 poses, both with -5.6 kcal/mol affinity and are shown in Supplementary Figure 14 and ranked by binding affinity in Figure 6. The docking results of the *CCT3* protein with ICG 001 resulted in 2 poses. The first pose had an affinity of -9.3 kcal/mol while the second pose had an affinity of -8.9 kcal/mol. The results were ranked by binding affinity and visualized in Figure 6.

**Figure 5.**
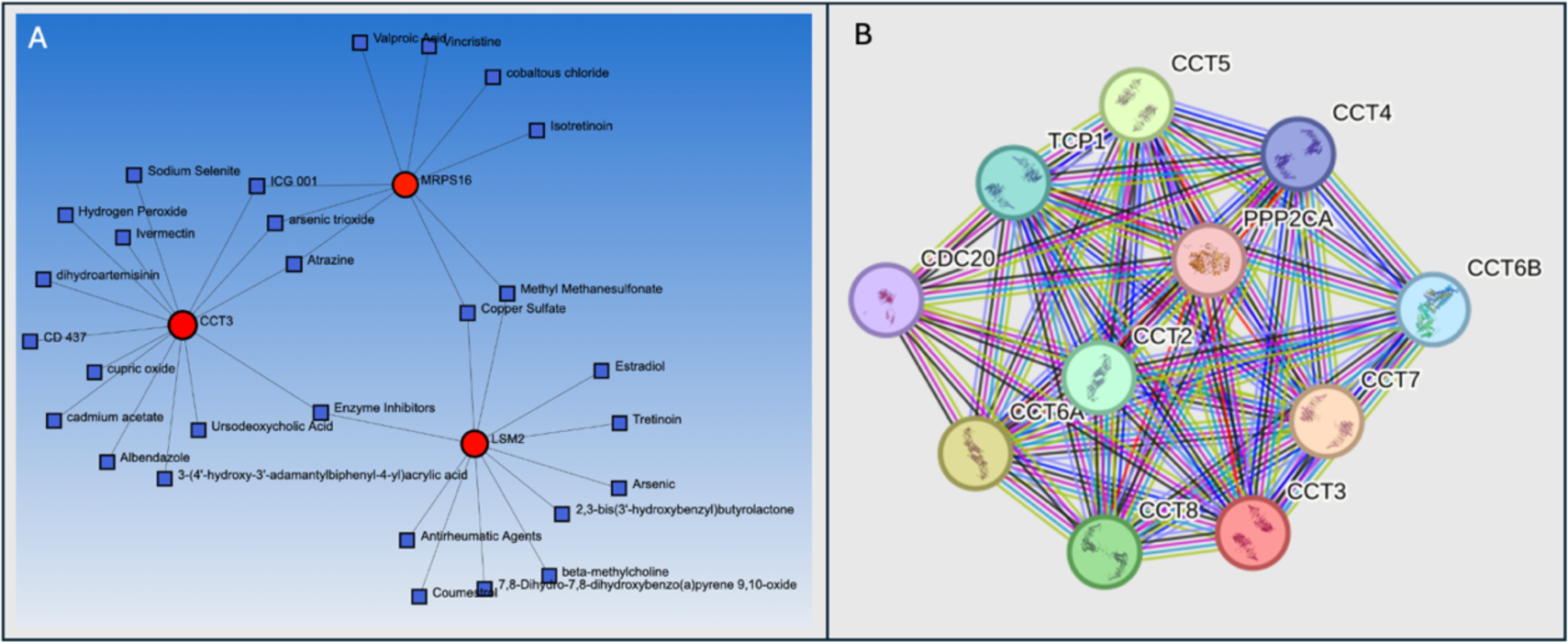
Identification of ligands for molecular docking and protein-protein interactions. (A) Node results from Network Analyst search of *CCT3*, *LSM2*, and *MRPS16*. Red circles indicate the target genes and blue squares denote the potential ligands. (B) Highly connective sub-network of relevant node cluster from STRING protein-protein interaction (PPI) for *CCT3*. The module consists of 10 other hub genes with a high degree of connectivity (degree > 50): *CCT5, TCP1, CDC20, CCT4, PPP2CA, CCT2, CCT6A, CCT6B, CCT7,* and *CCT8*.

**Figure 6.**
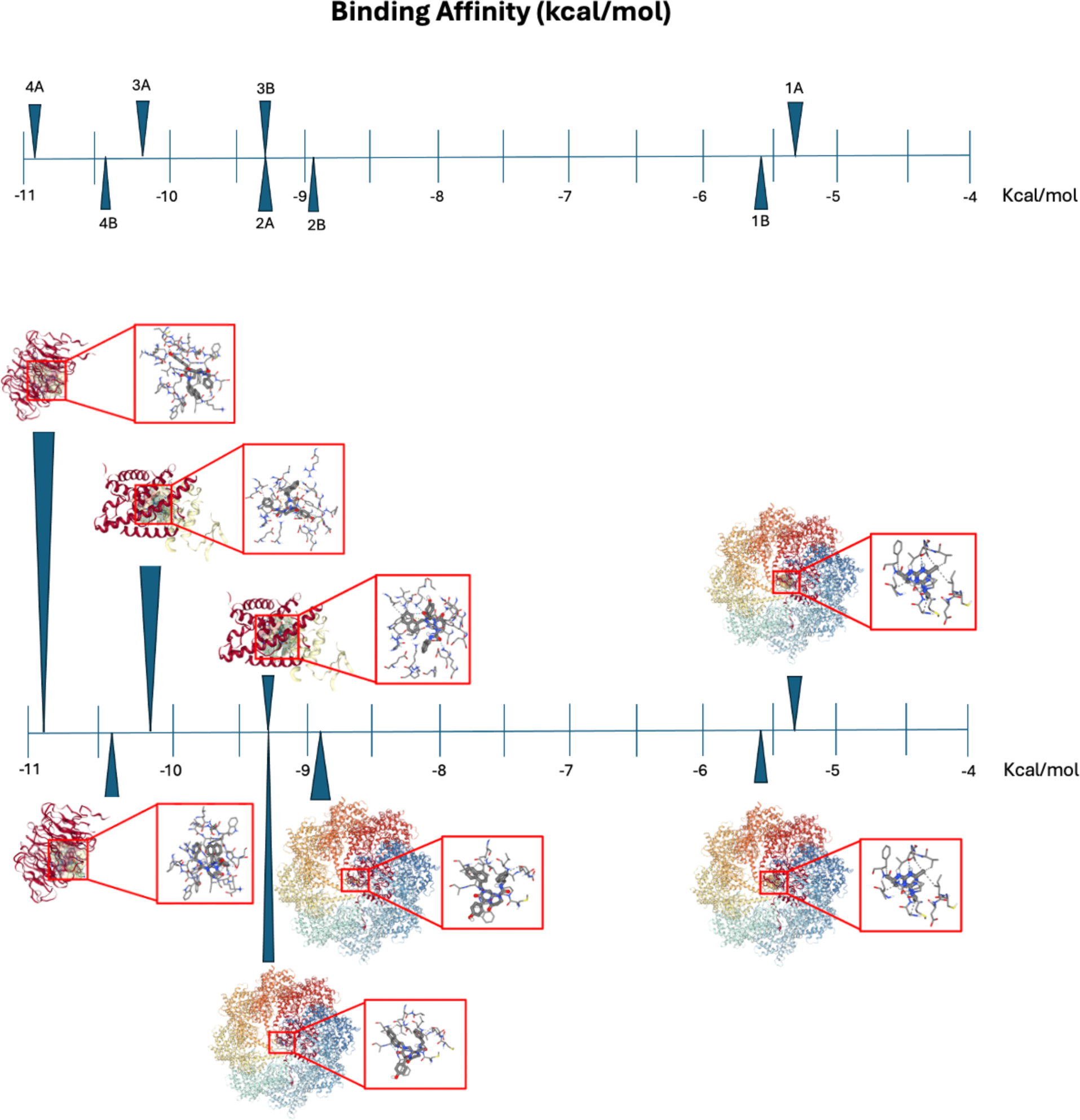
Molecular docking results. Seamdock results showing the ordered (1A) molecular docking position of Altrazine (PubChem CID 2256) in 7TTT for Pose 1 with binding affinity of -5.3 kcal/mol, (1B) molecular docking position of Altrazine (PubChem CID 2256) in 7TTT (*CCT3* protein) for Pose 2 with binding affinity of -5.6 kcal/mol, (2A) molecular docking position of ICG 001 (PubChem CID 11238147) in 7TTT (*CCT3* protein) for Pose 1 with binding affinity of -9.3 kcal/mol, and (2B) molecular docking position of ICG 001 (PubChem CID 11238147) in 7TTT (*CCT3* protein) for Pose 2 with binding affinity of -8.9 kcal/mol. (3A) molecular docking position of ICG001 (PubChem CID 11238147) docked to 4IYP (PPP2CA protein) in Pose 1 with binding affinity of -10.2 kcal/mol. (3B) molecular docking position of ICG001 (PubChem CID 11238147) docked to 4IYP (PPP2CA protein) in Pose 2 with binding affinity of -9.3 kcal/mol. (4A) molecular docking position of ICG001 (PubChem CID 11238147) docked to 4GGC (*CDC20* protein) in Pose 1 with binding affinity of -10.9 kcal/mol (4B) molecular docking position of ICG001 (PubChem CID 11238147) docked to 4GGC (*CDC20* protein) in Pose 2 with binding affinity of -10.4 kcal/mol.

### STRING analysis and Molecular Docking

The STRING database identified a PPI network of 11 genes as predicted functional partners to *CCT3*. Of those 11 genes, 9 were from the *CCT* family (*CCT4, CCT5, CCT2, CCT7, CCT8, CCT6A, CCT6B, TCP1*) and 2 were separate from it (*PPP2CA* and *CDC20*), as shown in Figure 5B. The molecular docking results of the *PPP2CA* protein with ICG 001 resulted in 2 poses: the first with binding affinity of -10.2 kcal/mol as shown in Figure 6.3A and the second with binding affinity of-9.3 kcal/mol shown in Figure 6.3B. The molecular docking results of *CDC20* with ICG001 also resulted in 2 poses: the first pose had the greatest binding affinity of -10.9 kcal/mol shown in Figure 6.4A and the second pose had binding affinity of -10.4 kcal/mol shown in Figure 6.4B.

### Functional and pathway enrichment analysis

To determine the molecular mechanisms underlying the ethnic disparities observed in distinct biological differences of TNBC between African American and Caucasian patients, functional annotation and pathway enrichment analyses of the DEGs was performed using DAVID. To minimize redundancy, The gene query was limited to DEGs with an absolute log2 fold change (log2FC) greater than 1 (|log2FC| > 1 or |FC| > 2.0). A collection of top ontology terms with significantly enriched biological processes, cellular components, and molecular functions are included in Supplementary Table 1.

The results analysis indicated that significant pathway overrepresentation primarily related to the upregulated genes in both datasets. As shown in Figure 3 and Table 3, the upregulated genes in GSE37751 were heavily focused on biological processes: cell division (GO:0051301), DNA repair (GO:0006281), chromosome segregation (GO:0007059), cell cycle (GO:0007049), mitotic sister chromatid segregation (GO:0000070), and DNA replication (GO:0006260). The upregulated genes were significantly concentrated in cellular components relevant to the nucleus, including nucleoplasm, centrosome, kinetochore, and chromosome. Enriched molecular functions included protein binding (GO:0005515), RNA binding (GO:0003723), ATP binding (GO:0005524), and ATPase activity (GO:0016887).

**Table 3.**
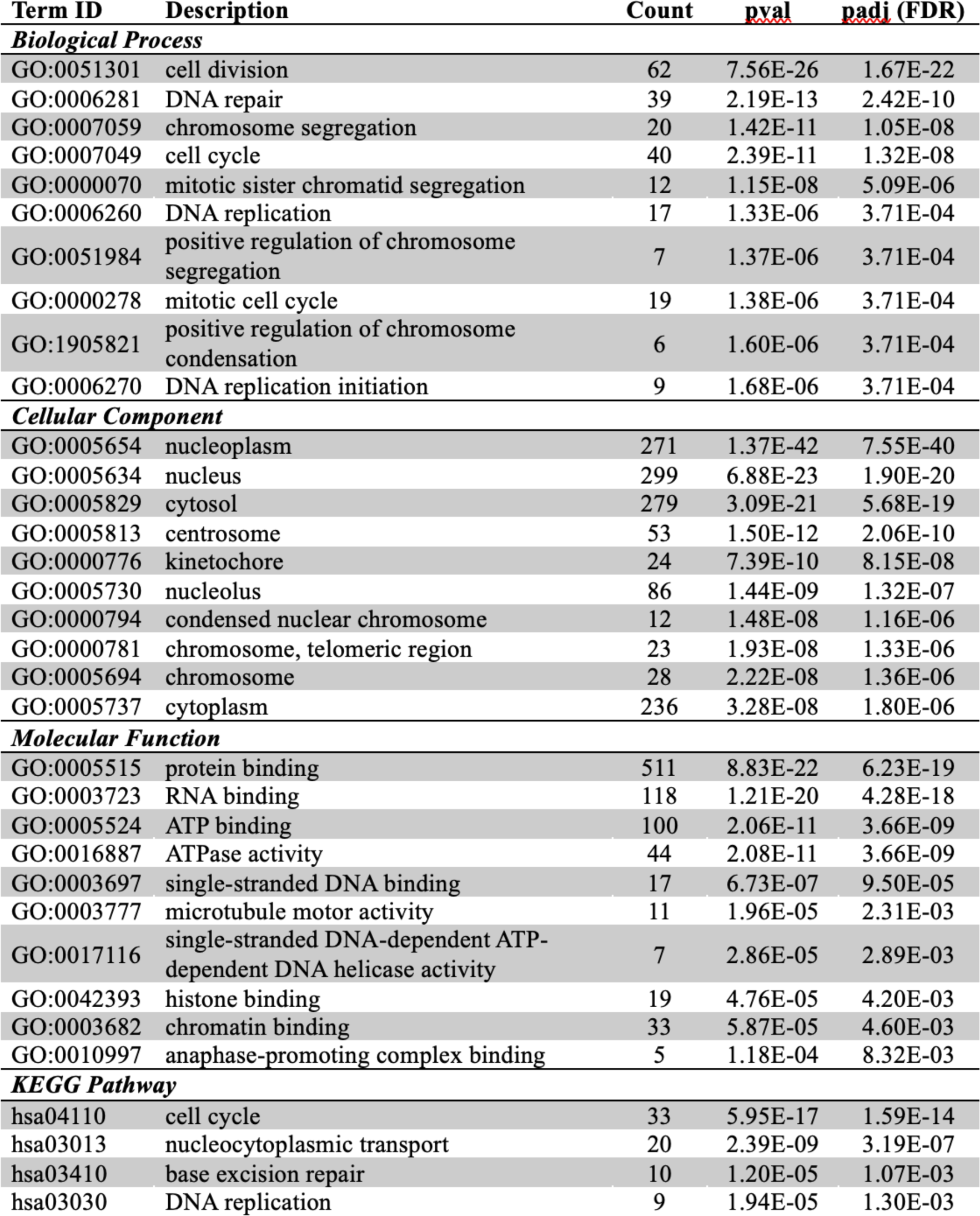
Gene Ontology (GO) and KEGG enrichment analysis for DEGs from GSE37751. The four major areas of the table enrichment analysis include biological process, cellular component, molecular function, and KEGG pathway. For each area, the Term ID, term description, count, p-value (pval), and adjusted p-value (padj (FDR)) are provided.

| Term ID | Description | Count | pval | padj (FDR) |
| --- | --- | --- | --- | --- |
| **Biological Process** |  |  |  |  |
| GO:0051301 | cell division | 62 | 7.56E-26 | 1.67E-22 |
| GO:0006281 | DNA repair | 39 | 2.19E-13 | 2.42E-10 |
| GO:0007059 | chromosome segregation | 20 | 1.42E-11 | 1.05E-08 |
| GO:0007049 | cell cycle | 40 | 2.39E-11 | 1.32E-08 |
| GO:0000070 | mitotic sister chromatid segregation | 12 | 1.15E-08 | 5.09E-06 |
| GO:0006260 | DNA replication | 17 | 1.33E-06 | 3.71E-04 |
| GO:0051984 | positive regulation of chromosome segregation | 7 | 1.37E-06 | 3.71E-04 |
| GO:0000278 | mitotic cell cycle | 19 | 1.38E-06 | 3.71E-04 |
| GO:1905821 | positive regulation of chromosome condensation | 6 | 1.60E-06 | 3.71E-04 |
| GO:0006270 | DNA replication initiation | 9 | 1.68E-06 | 3.71E-04 |
| **Cellular Component** |  |  |  |  |
| GO:0005654 | nucleoplasm | 271 | 1.37E-42 | 7.55E-40 |
| GO:0005634 | nucleus | 299 | 6.88E-23 | 1.90E-20 |
| GO:0005829 | cytosol | 279 | 3.09E-21 | 5.68E-19 |
| GO:0005813 | centrosome | 53 | 1.50E-12 | 2.06E-10 |
| GO:0000776 | kinetochore | 24 | 7.39E-10 | 8.15E-08 |
| GO:0005730 | nucleolus | 86 | 1.44E-09 | 1.32E-07 |
| GO:0000794 | condensed nuclear chromosome | 12 | 1.48E-08 | 1.16E-06 |
| GO:0000781 | chromosome, telomeric region | 23 | 1.93E-08 | 1.33E-06 |
| GO:0005694 | chromosome | 28 | 2.22E-08 | 1.36E-06 |
| GO:0005737 | cytoplasm | 236 | 3.28E-08 | 1.80E-06 |
| **Molecular Function** |  |  |  |  |
| GO:0005515 | protein binding | 511 | 8.83E-22 | 6.23E-19 |
| GO:0003723 | RNA binding | 118 | 1.21E-20 | 4.28E-18 |
| GO:0005524 | ATP binding | 100 | 2.06E-11 | 3.66E-09 |
| GO:0016887 | ATPase activity | 44 | 2.08E-11 | 3.66E-09 |
| GO:0003697 | single-stranded DNA binding | 17 | 6.73E-07 | 9.50E-05 |
| GO:0003777 | microtubule motor activity | 11 | 1.96E-05 | 2.31E-03 |
| GO:0017116 | single-stranded DNA-dependent ATP-dependent DNA helicase activity | 7 | 2.86E-05 | 2.89E-03 |
| GO:0042393 | histone binding | 19 | 4.76E-05 | 4.20E-03 |
| GO:0003682 | chromatin binding | 33 | 5.87E-05 | 4.60E-03 |
| GO:0010997 | anaphase-promoting complex binding | 5 | 1.18E-04 | 8.32E-03 |
| **KEGG Pathway** |  |  |  |  |
| hsa04110 | cell cycle | 33 | 5.95E-17 | 1.59E-14 |
| hsa03013 | nucleocytoplasmic transport | 20 | 2.39E-09 | 3.19E-07 |
| hsa03410 | base excision repair | 10 | 1.20E-05 | 1.07E-03 |
| hsa03030 | DNA replication | 9 | 1.94E-05 | 1.30E-03 |

Furthermore, the upregulated genes in GSE142731 were involved in biological processes, including telomere organization (GO:0032200), nucleosome assembly (GO:0006334), protein localization to CENP-A containing chromatin (GO:0061644), DNA replication-dependent nucleosome assembly (GO:0006335), and heterochromatin assembly (GO:0031507). Consistent with the results for GSE37751, the upregulated genes for GS142731 shown in Table 4 were heavily enriched in cellular components pertaining to the nucleus, including: nucleosome, nuclear chromosome, nucleoplasm, and CENP-A containing nucleosome. Associated molecular function categories included structural constituent of chromatin (GO:0030527), protein heterodimerization (GO:0046982), as well as protein binding (GO:0005515) and RNA binding (GO:0003723). The downregulated DEGs only exhibited enrichment in extracellular matrix cellular component category.

**Table 4.**
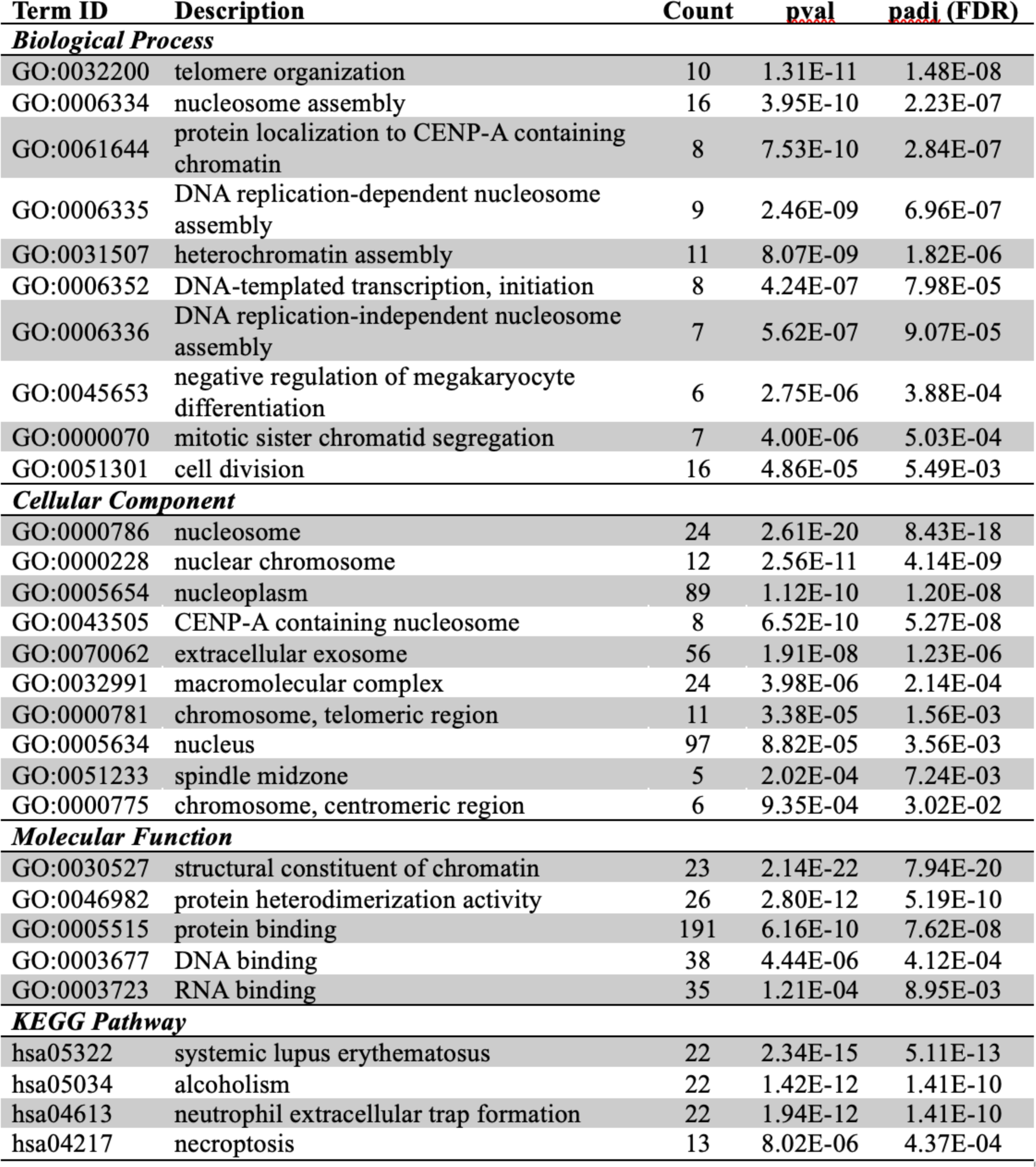
Gene Ontology (GO) and KEGG enrichment analysis for DEGs from GSE142731. The four major areas of the table enrichment analysis include biological process, cellular component, molecular function, and KEGG pathway. For each area, the Term ID, term description, count, p-value (pval), and adjusted p-value (padj (FDR)) are provided.

**Table 5.** Significance results for African American and European American patients with triple negative breast cancer. DEGs include *CCT3*, *LSM2*, and *MRPS16* with corresponding log2 fold change (log2FC) and p-values (p-val) for GSE37751 and GSE14731.

| Entrez ID | Gene | GSE37751 |  | GSE142731 |  | Description |
| --- | --- | --- | --- | --- | --- | --- |
|  |  | log2FC | pval | log2FC | pval |  |
| 7203 | *CCT3* | 0.452 | 3.01E-03 | 0.826 | 2.25E-05 | chaperonin containing TCP1 subunit 3 |
| 57819 | *LSM2* | 1.28 | 1.51E-05 | 0.645 | 3.32E-04 | LSM2 homolog, U6 small nuclear RNA and mRNA degradation associated |
| 51021 | *MRPS16* | 0.426 | 3.41E-03 | 0.588 | 9.68E-05 | mitochondrial ribosomal protein S16 |

**Table 6.** Potential Ligands identified from Network Analyst input of *CCT3, LSM2*, and *MRPS16* search. Results include ID, Degree, and Betweenness for each Label.

| ID | Label | Degree | Betweenness |
| --- | --- | --- | --- |
| 7203 | CCT3 | 14 | 261.5 |
| 57819 | LSM2 | 11 | 214 |
| 51021 | MRPS16 | 9 | 152.5 |
| D004791 | Enzyme Inhibitors | 2 | 113.5 |
| D019327 | Copper Sulfate | 2 | 32.5 |
| D008741 | Methyl Methanesulfonate | 2 | 32.5 |
| C006632 | arsenic trioxide | 2 | 24.83 |
| D001280 | Atrazine | 2 | 24.83 |
| C492448 | ICG 001 | 2 | 24.83 |
| C029497 | 2,3-bis(3'-hydroxybenzyl)butyrolactone | 1 | 0 |
| C472791 | 3-(4'-hydroxy-3'-adamantylbiphenyl-4-yl)acrylic acid | 1 | 0 |
| D015123 | 7,8-Dihydro-7,8-dihydroxybenzo(a)pyrene 9,10-oxide | 1 | 0 |
| D015766 | Albendazole | 1 | 0 |
| D018501 | Antirheumatic Agents | 1 | 0 |
| D001151 | Arsenic | 1 | 0 |
| C044887 | beta-methylcholine | 1 | 0 |
| C028031 | cadmium acetate | 1 | 0 |
| C099555 | CD 437 | 1 | 0 |
| C018021 | cobaltous chloride | 1 | 0 |
| D003375 | Coumestrol | 1 | 0 |
| C030973 | cupric oxide | 1 | 0 |
| C039060 | dihydroartemisinin | 1 | 0 |
| D004958 | Estradiol | 1 | 0 |
| D006861 | Hydrogen Peroxide | 1 | 0 |
| D015474 | Isotretinoin | 1 | 0 |
| D007559 | Ivermectin | 1 | 0 |
| D018038 | Sodium Selenite | 1 | 0 |
| D014212 | Tretinoin | 1 | 0 |
| D014580 | Ursodeoxycholic Acid | 1 | 0 |
| D014635 | Valproic Acid | 1 | 0 |
| D014750 | Vincristine | 1 | 0 |

## Discussion

TNBC differs from other classifications of breast cancer in its metastatic potential [22] and rapidly diminishing survival rate as the cancer progresses. A study predicted that over 50% of TNBC patients will develop distant metastasis [23] in which the cancer will have spread to distant locations in the body, such as other organs. TNBC tumors are notably less responsive to traditional chemotherapy and more difficult to treat due to a lack of targetable receptors. The present study aims to identify biomarkers that may be further examined as novel strategies for treatment of TNBC in African American patients.

The Warburg Effect refers to a phenomenon observed in oncogenesis that involves metabolic reprogramming to support uncontrolled proliferation and tumor survival and is often described as the hallmark of cancer cells [24]. In this process, tumor cells exploit mitochondrial respiration by upregulating aerobic glycolysis for rapid ATP production rather than oxidative phosphorylation. Aerobic glycolysis is favored by cancer cells to facilitate the elevated rate of glucose consumption and subsequent production of lactate even under normal oxygen concentrations.

The preference for aerobic glycolysis is not only an inefficient alternative for generating ATP but also impedes proper execution of downstream cell signaling mechanisms to support metabolic reprogramming [25]. While the biological implications of the Warburg Effect in carcinogenesis is still an active area of research, a major focus lies on the effects of endogenous lactate as an oncometabolite, which stands out as a critical concern. Numerous studies demonstrate that lactate acts as a regulator of various cellular processes involved in cancer progression. Aberrant accumulation of the metabolite is reported to induce a variety of metabolic alterations that may contribute to the aggressive phenotype and metastatic capabilities of TNBC cells. In the current study, gene expression data of breast cancer patients was downloaded and analyzed to identify genes that are differentially expressed between African American and Caucasian women with TNBC.

The three survival-related genes, *CCT3*, *LSM2*, and *MRPS16*, showed correlation with shorter survival times in AA patients. The three genes showed significantly differential expression among African American and European American patients with triple negative breast cancer and have roles in significantly enriched cellular processes, which could contribute to the aggressive phenotypes and poor prognosis of TNBC in African American patients. The survival analysis, and differential expression results show that *CCT3, LSM2*, and *MRPS16* could be potential diagnostic and prognostic biomarkers for triple negative breast cancer.

To explore potential binding targets for the genes of interest (*CCT3, LSM2,* and *MRPS16*), meta-analysis was performed using the Network Analyst server. Atrazine and ICG 001 were identified as potential ligands, both with 2 degrees (connecting to *CCT3* and *MPRS16*), a Betweenness score of 24.8333, and molecular docking results with the *CCT3* protein resulted in strong binding affinities. However, Atrazine is an herbicide and therefore a health hazard, so it is likely not optimal for targeting the *CCT3* protein in homo sapiens. ICG 001 has a role in Wnt/beta-catenin pathways without the health hazards of Atrazine and therefore may be a more optimal option for targeting *CCT3* proteins. While arsenic trioxide was identified as a potential binding target for *CCT3* and *MRPS16*, copper sulfate and methyl methanosulfonate as potential ligands for *MRPS16* and *LSM2*, and enzyme inhibitors as ligands for *CCT3* and *LSM2*, these were deemed unsuitable for attempting molecular docking due to the lack of a 3D structure (like arsenic trioxide and copper sulfate) or broad nature (like enzyme inhibitors). The remaining ligands were identified but not used for molecular binding as *LSM2* and *MRPS16* did not have a protein structure derived from homo sapiens and therefore were not useful for clinical investigation. Therefore, this study continued investigating *CCT3* as a potential biomarker.

The STRING database identified *PPP2CA* and *CDC20* as distinct predicted functional partners to *CCT3*. The molecular docking results of *CDC20* with ICG001 also resulted in 2 poses: the first pose had the greatest binding affinity of - 10.9 kcal/mol shown in Figure 6.4A and the second pose had second highest binding affinity of -10.4 kcal/mol. The molecular docking results of the *PPP2CA* protein with ICG 001 resulted in 2 poses: the first with the next highest binding affinity of -10.2 kcal/mol and the second with binding affinity of-9.3 kcal/mol. However, having investigated the survival analysis of *CDC20* as a differentially expressed gene in the dataset, as shown in Supplementary Figure 13, and the TCGA expression profile in cancer cell lines, as shown in Supplementary Figure 10 with significant expression in primary tumor, Triple Negative samples, African-American populations, and TP-53 mutant.

Overall, pathway analysis of the upregulated DEGs from datasets GSE37751 and GSE142731 demonstrated significant enrichment in GO terms relevant to cell division, mitotic sister chromatid segregation, nucleoplasm, nucleus, chromosome (telomeric region), protein binding, and RNA binding. KEGG analysis showed overrepresentation of pathways that may play a role in breast cancer development, including systemic lupus erythematosus, alcoholism, cell cycle, and necroptosis for GSE142731. The analysis for GSE37751 showed overrepresentation of pathways that could have a role in breast cancer development, including cell cycle, nucleocytoplasmic transport, base excision repair, DNA replication, amyotrophic lateral sclerosis, and oocyte meiosis. A protein interaction network was generated to visualize the roles and relationships between the DEGs. *CCT3* and *CDC20*, with significant protein-protein interactions with promising TCGA or survival results, and effective molecular docking, could serve as potential therapeutic targets.

## Conclusion

Data from public breast cancer cohorts was used to perform a bioinformatic analysis and identify expression patterns associated with poor survival outcomes of African American patients with TNBC. Our findings support the notion that racial disparities between African American and Caucasian patients with TNBC exist at the molecular level. Evidence suggests the derived genes (*CCT3*, *LSM2*, and *MRPS16*) could serve as candidate biomarkers for diagnosis, prognosis, and novel therapeutic targets for TNBC in African American patients. Molecular docking methods identified ICG001 as a potential therapeutic target for binding the prospective protein biomarker, *CCT3*. Further research on the clinical implications of genes *CCT3*, *LSM2*, and *MRPS16,* their molecular targets, and protein-protein interactions could provide new insights into the potential relationship of these genes with outcomes of TNBC.

## Materials and Methods

### Methodology Pipeline

The methodology pipeline used throughout is pictured in Figure 7.

**Fig 7.**
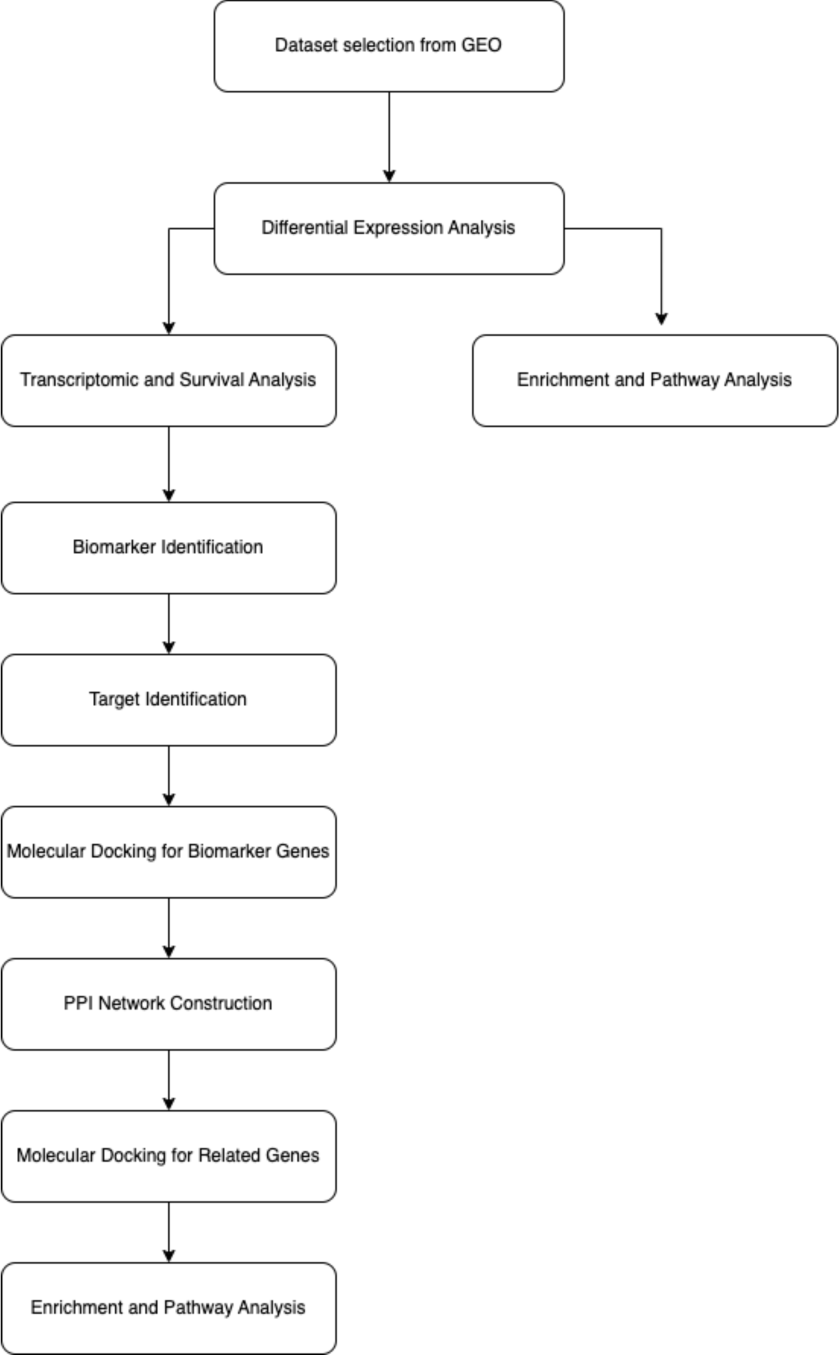
Methods Pipeline. Methods include Dataset selection from GEO, differential expression, transcriptomic and survival analysis, enrichment and pathway analysis, biomarker identification, target identification, molecular docking for biomarker genes, PPI network construction, molecular docking for related genes, enrichment and pathway analysis.

### Datasets collection

Datasets selected in the study were downloaded from the Gene Expression Omnibus (GEO) database repository (https://www.ncbi.nlm.nih.gov/geo/) (accessed on 10 March 2023) [16], containing publicly available genomic data from human breast tissue samples. The following criteria was used to select the datasets (i) Organism - *Homo sapiens*, (ii) Datatype - expression profiling, (iii) breast cancer subtype classification - triple-negative breast cancer, and (iv) patient ethnicity - African American and European American patients. Analysis of expression data from the GEO datasets were based on GPL6244 (Affymetrix Human Gene 1.0 ST Array) and GPL16791 (Illumina HiSeq 2500) platforms, respectively.

### Differential expression analysis

Differentially expressed genes (DEGs) were detected using GEO2R, an R-based analytic tool [17](https://www.ncbi.nlm.nih.gov/geo/geo2r/) (accessed on 10 March 2023), among the two designated sample groups: African American patients with TNBC, European American patients with TNBC. Differential expression was defined based on adjusted P value cutoff (padj < 0.05), calculated using Benjamini–Hochberg (FDR) method. No restrictions were applied to log2 fold change (log2FC) threshold. DEG screening was performed on September 22, 2022.

### Transcriptomic and survival analysis

To identify candidate genetic markers associated with poorer prognoses in TNBC patients of African American descent, a survival analysis was conducted utilizing the University of Alabama Cancer portal (UALCAN) database [20, 21] (www.ualcan.path.uab.edu/index.html) (accessed on 10 March 2023). The research assessed transcriptomic data from TCGA of the DEGs for significant differential expression between breast invasive carcinoma tumor cells and healthy cells and correlation with TNBC-specific survival probabilities that vary based on patient ethnicity.

### Target Identification

To identify potential ligands for molecular docking, meta-analysis was performed for the genes *CCT3, LSM2*, and *MRPS16* using Network Analyst [28, 29, 30, 31, 32] (https://www.networkanalyst.ca/) (accessed on 20 April 2024), a web-based server. The study used the ligands of interest for molecular docking with the gene products. The 3D structures for the ligands and receptors were downloaded from RCSB PDB [33] (https://www.rcsb.org/) (accessed on 20 April 2024) and PubChem [34] (https://pubchem.ncbi.nlm.nih.gov/) (accessed on 20 April 2024).

### Molecular Docking

The ligands were prepared as PDB files using PyMol [35] (https://pymol.org/)(accessed) on 21 April 2024), an open-source visualization tool. Molecular docking was performed using SeamDock [26, 27] (https://bioserv.rpbs.univ-paris-diderot.fr/services/SeamDock/) (accessed on 21 April 2024), a web-based molecular docking server. The molecular docking results were assessed for effective binding interactions of the receptor-ligand complex. Parameters for molecular docking the Altrazine ligand into the *CCT3* gene protein with PDB ID 7TTT from SeamDock included the software of Vina, Spacing = 1, Mode number = 8, Energy Range = 0.1, and Exhaustivness = 8. Parameters for molecular docking the ICG 001 ligand into the *CCT3* gene protein (PDB ID 7TTT), into the *CDC20* gene protein (PDB ID 4GGC) and into the *PPP2AC* gene protein (PDB ID 4IYP) in SeamDock included the software of Vina, Spacing = 1, Mode number = 2, Energy Range = 5, and Exhaustivness = 8.

### PPI Network Construction and Identification of Co-expression Modules

To extract significant co-expression modules and further explore functional relationships among the DEGs, a protein-protein interaction (PPI) network was constructed using STRING [19] (https://string-db.org/) (accessed on 1 May 2024), an online database of known and predicted protein-protein interactions.

### Gene ontology (GO) and Kyoto encyclopedia of genes and genomes (KEGG) pathway enrichment analysis

The DEGs were screened for enriched biological pathways and molecular functions using the Database for Annotation, Visualization, and Integrated Discovery (DAVID) functional annotation tool, Enrichr, and Metascape [18](www.metascape.org) (accessed on 10 March 2023), a web-based bioinformatics tool.

## Acknowledgments Funding

National Institute on Minority Health and Health Disparities Award U54MD015946

## Author contributions

Conceptualization: KNC, HAM, JR, SY

Experimental recordings: KNC, HAM, JR, SY

Data analysis: KNC, HAM, JR, SY

Computational modeling: KNC, HAM, JR, SY

Figure creation: KNC, HAM, JR, SY

Writing: KNC, HAM, JR, SY

## Data and materials availability

All data are available in the main text or the supplementary materials.

## Supplementary Materials

Supplementary Materials include Supplementary Figures, and Supplementary Tables.

## Supplementary Materials

### Supplementary Figures

**Figure S1.**
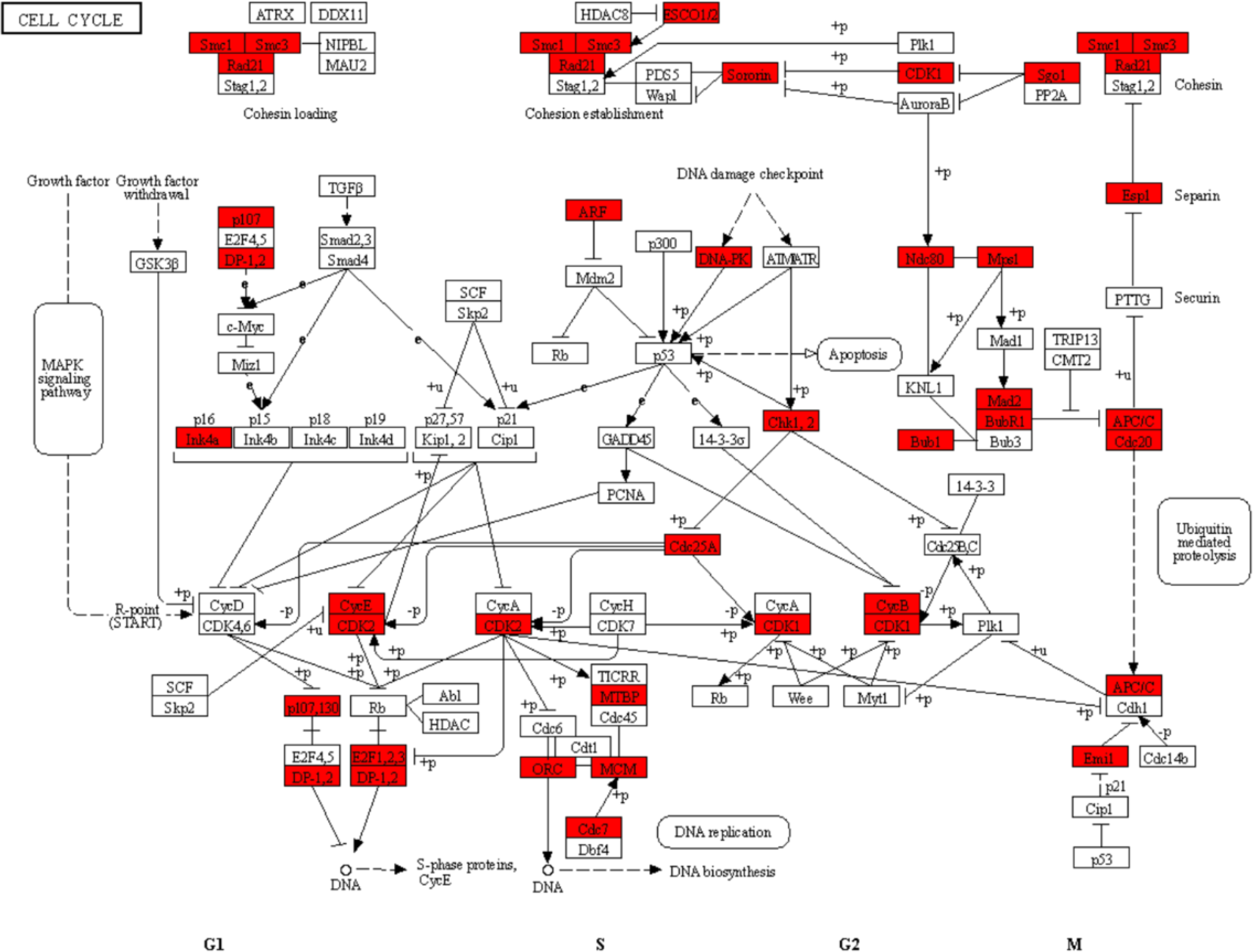
KEGG Pathway: Cell cycle (hsa04110) with annotations of upregulated DEGs. The cell cycle pathway includes four main phases: Gap 1 (G1), DNA replication synthesis (S), Gap 2 (G2), and Mitosis (M). Related pathways include apoptosis, MAPK signaling pathway, DNA replication, and Ubiquitin mediated proteolysis. Some of the upregulated genes include cyclin-dependent kinases (CDK) which act as regulatory enzymes and target specific transcription factors (E2F), structural maintenance of chromosomes (Smc), and Chk1,2 involved with tumor suppression, among others highlighted in red (upregulated).

**Figure S2.**
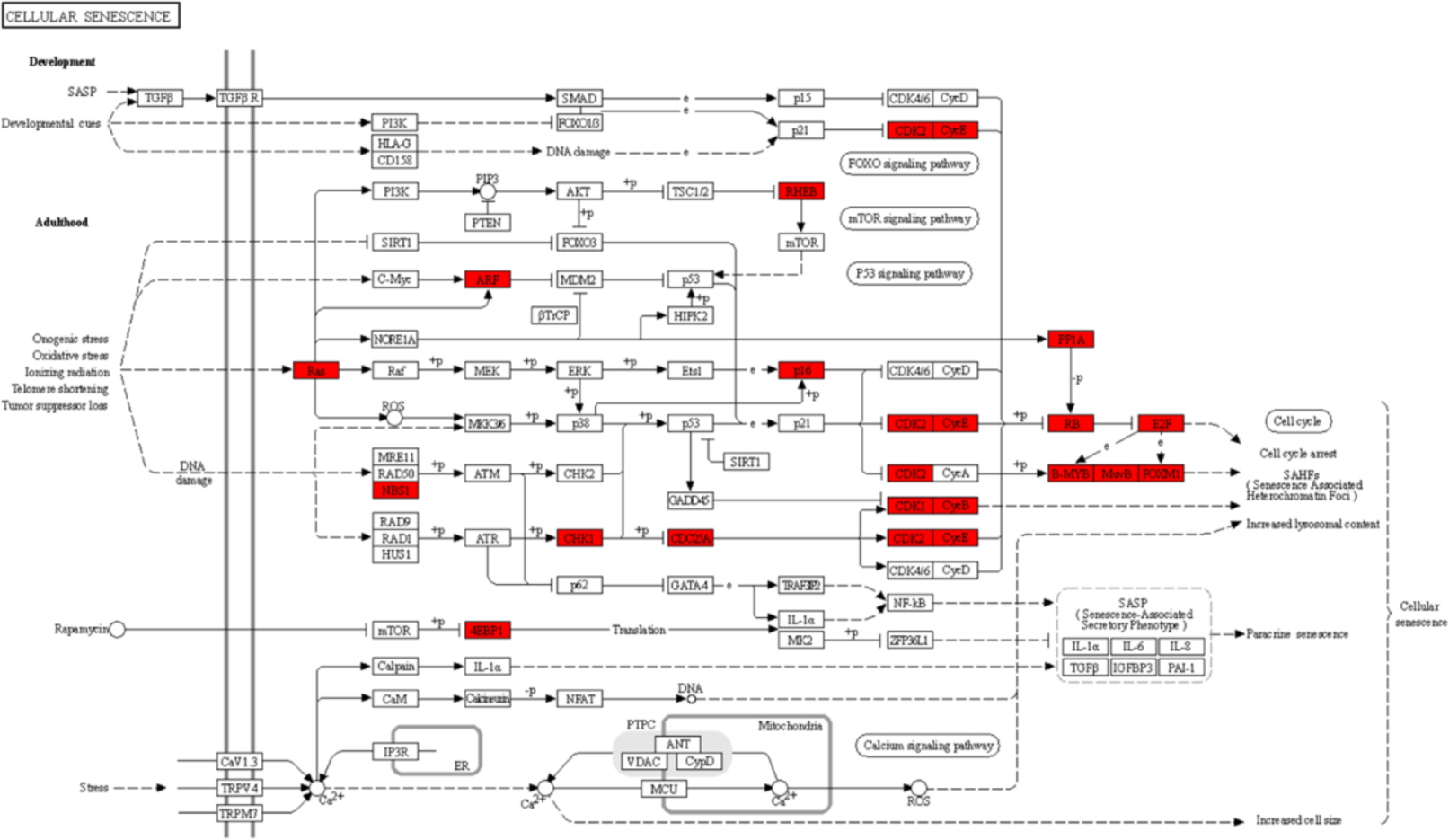
KEGG Pathway: Cellular senescence (hsa04218) with annotations of upregulated DEGs. Cellular senescence is the process of irreversible cellular arrest. The pathway is related to the calcium signaling, FoxO signaling, cell cycle, p53 signaling, and mTOR signaling pathways. Genes upregulated (highlighted red) in cellular senescence include Ras-related 2 (Ras), cyclin dependent kinase inhibitor 2A (ARF), nibrin (NBS1), eukaryotic translation initiation factor 4E binding protein 1 (4EBP1), checkpoint kinase 1 (CHK1), M-phase inducer phosphatase 1 (CDC25A), Ras homolog enriched in brain (RHEB), cyclin dependent kinase inhibitor 2A (p16), cyclin dependent kinase 1 & 2 (CDK1 & 2), G2/mitotic-specific cyclin-B3 & G1/S-specific cyclin-E1 (CycB & E), protein phosphatase 1 catalytic subunit alpha (PP1A), retinoblastoma-associated protein (RB), transcription factor E2F1 (E2F), MYB proto-oncogene like 2 (B-MYB), lin-54 DREAM MuvB core complex component (MuvB), and forkhead box M1 (FOXM1).

**Figure S3.**
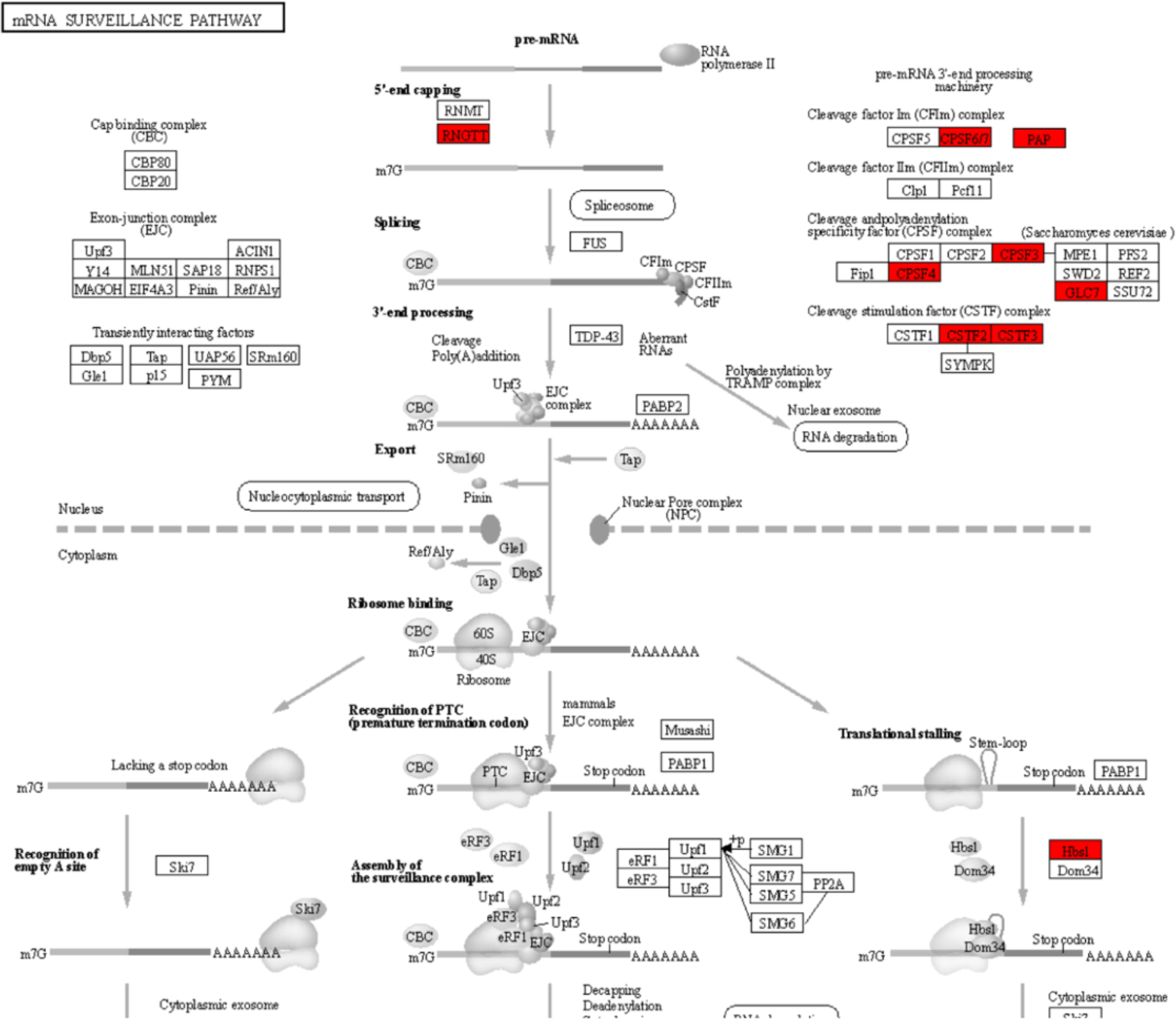
KEGG Pathway: mRNA surveillance (hsa03015) with annotations of upregulated DEGs. The mRNA surveillance pathway is primarily involved in quality control for detection and degradation of abnormal mRNA. Related pathways include nucleocytoplasmic transport, RNA degradation, and spliceosome. The upregulated genes include RNA guanylyltransferase and 5’-phosphatase (RNGTT), cleavage and polyadenylation specific factor 6 & 7 (CPPSF6/7), poly(A) polymerase alpha (PAP), cleavage and polyadenylation specific factor 3 & 4 (CPSF3 & 4), protein phosphatase 1 catalytic subunit alpha (GLC7), cleavage stimulation factor subunit 2 & 3 (CSTF2 & 3), and HBS1 like translational GTPase (Hbs1).

**Figure S4.**
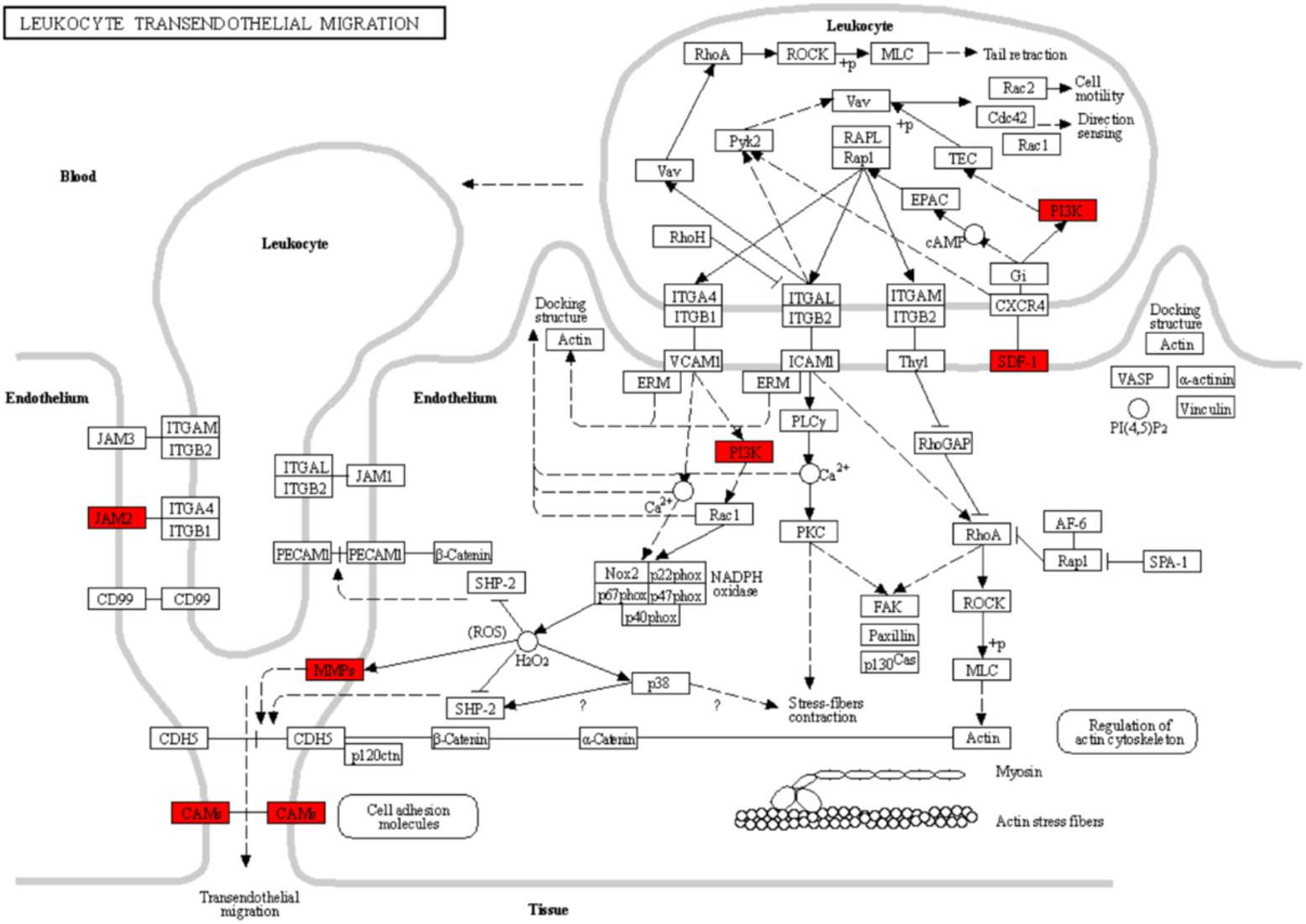
KEGG Pathway: Leukocyte transendothelial migration (hsa04670) with annotations of downregulated DEGs. The leukocyte transendothelial migration pathway is important for immune surveillance and inflammation. Related pathways include cell adhesion molecules and regulation of actin cytoskeleton. The downregulated genes (highlighted red) include junctional adhesion molecule 2 (JAM2), Cell adhesion molecules (CAMs), matrix metallopeptidase 2 (MMPs), phosphatidylinositol-4,5-bisphosphate 3-kinase catalytic subunit alpha (PI3K), and C-X-C motif chemokine ligand 12 (SDF-1).

**Figure S5.**
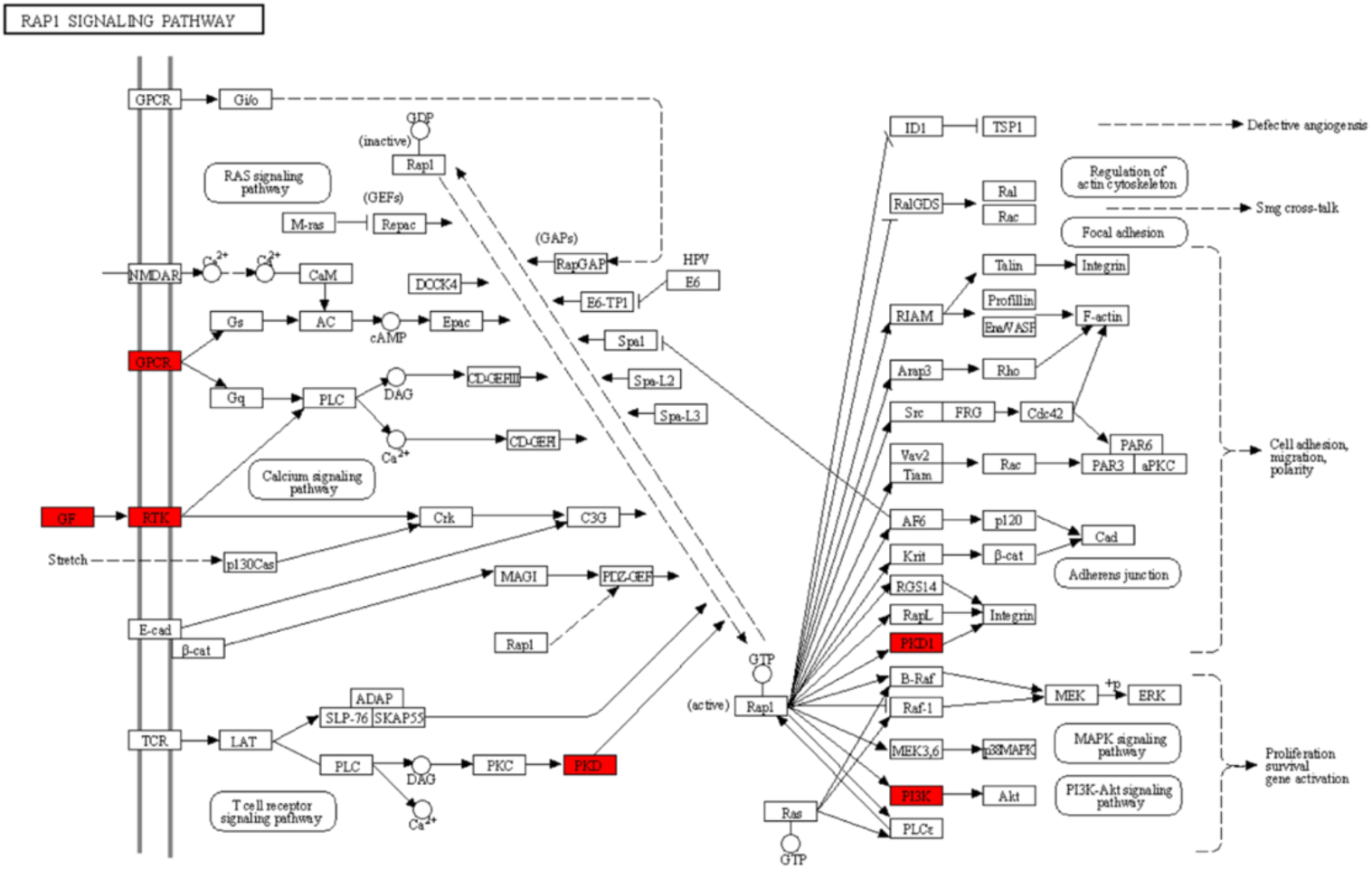
KEGG Pathway: Rap1 signaling (hsa04015) with annotations of downregulated DEGs. The RAP1 signaling pathway is an environmental information processing and signal transduction class that controls multiple diverse processes. It can result in defective angiogenesis, Smg cross-talk, cell adhesion, migration, and polarity, and proliferation survival gene activation. Related pathways include the RAS signaling pathway, T cell receptor signaling pathway, Regulation of actin cytoskeleton, focal adhesion, adherins junction, MAPK signaling pathway, and P13K signaling pathway. The downregulated genes (highlighted red) include Adenosine A2a receptor (GPCR), colony stimulating factor 1 (GF) and colony stimulating factor 1 receptor (RTK), protein kinase D (PKD and PKD1), and phosphatidylinositol-4,5-bisphosphate 3-kinase catalytic subunit alpha (PI3K).

**Figure S6.**
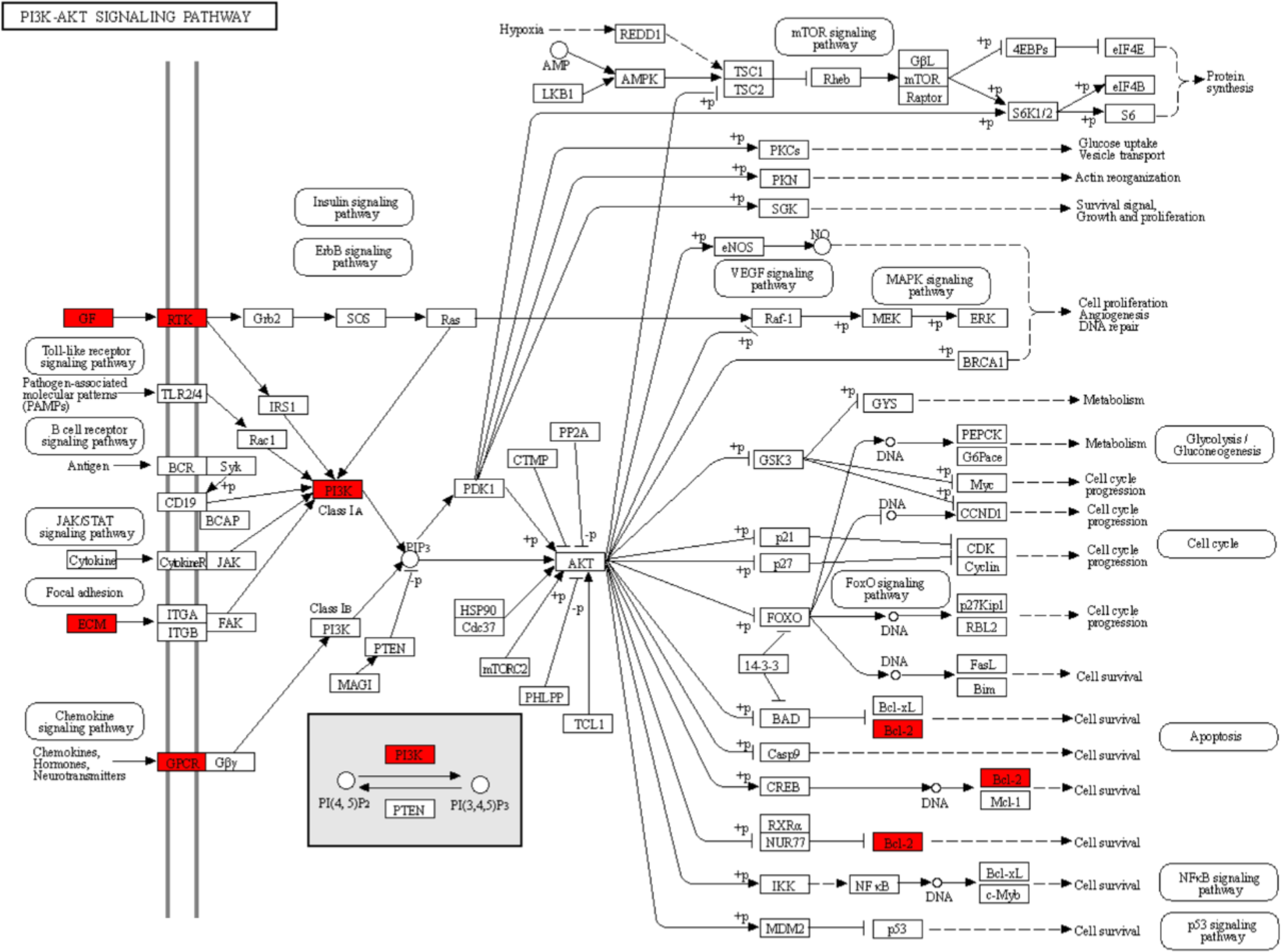
KEGG Pathway: PI3K-AKT signaling (hsa04151) with annotations of downregulated DEGs. The PI3K-AKT signaling pathway is important for apoptosis, protein synthesis, metabolism, and cell cycle. Related pathways include glycolysis/gluconeogenesis, MAPK signaling, ErB signaling, Chemokine signaling, NF-kappa B signaling, FoxO signaling, cell cycle, mTOR signaling, p53 signaling, apoptosis, VEGF signaling, focal adhesion, toll-like receptor signaling, JAK-STAT signaling, B cell receptor signaling, and insulin signaling. The downregulated genes (highlighted red) include colony stimulating factor 1 (GF), laminin subunit gamma 3 (ECM), lysophosphatidic acid receptor 6 (GPCR), colony stimulating factor 1 receptor (RTK), phosphatidylinositol-4,5-bisphosphate 3-kinase catalytic subunit alpha (PI3K), and BCL2 apoptosis regulator (Bcl-2).

**Figure S7.**
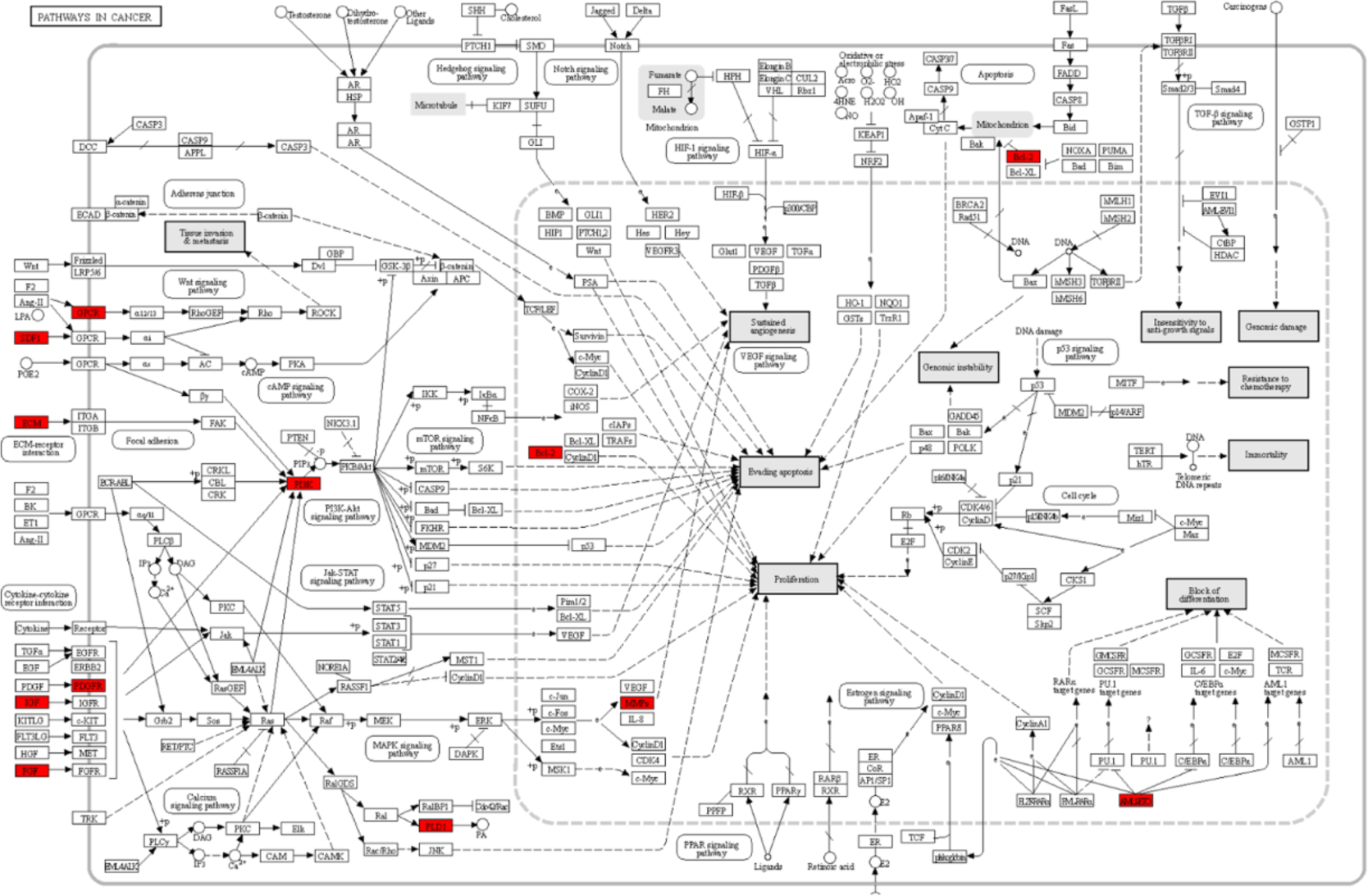
KEGG Pathway: Pathways in cancer (hsa05200) with annotations of downregulated DEGs. Pathways in cancer are related to PPAR signaling, MAPK signaling, Calcium signaling, cAMP signaling, cytokine-cytokine receptor interaction, HIF-1 signaling, cell cycle, p53 signaling, mTOR signaling, PI3K-Akt signaling, apoptosis, Wnt signaling, Notch signaling, Hedgehog signaling, TGF-beta signaling, VEGF signaling, focal adhesion, ECM-receptor interaction, JAK-STAT signaling, and estrogen signaling. Genes downregulated (highlighted red) include C-X-C motif chemokine ligand 12 (SDF1), laminin subunit gamma 3 (ECM), insulin like growth factor 1 (IGF), fibroblast growth factor 1 (FGF), phosphatidylinositol-4,5-bisphosphate 3-kinase catalytic subunit alpha (PI3K), phospholipase D1 (PLD1), BCL2 apoptosis regulator (Bcl-2), matrix metallopeptidase (MMPs), and RUNX family transcription factor 1.

**Figure S8.**
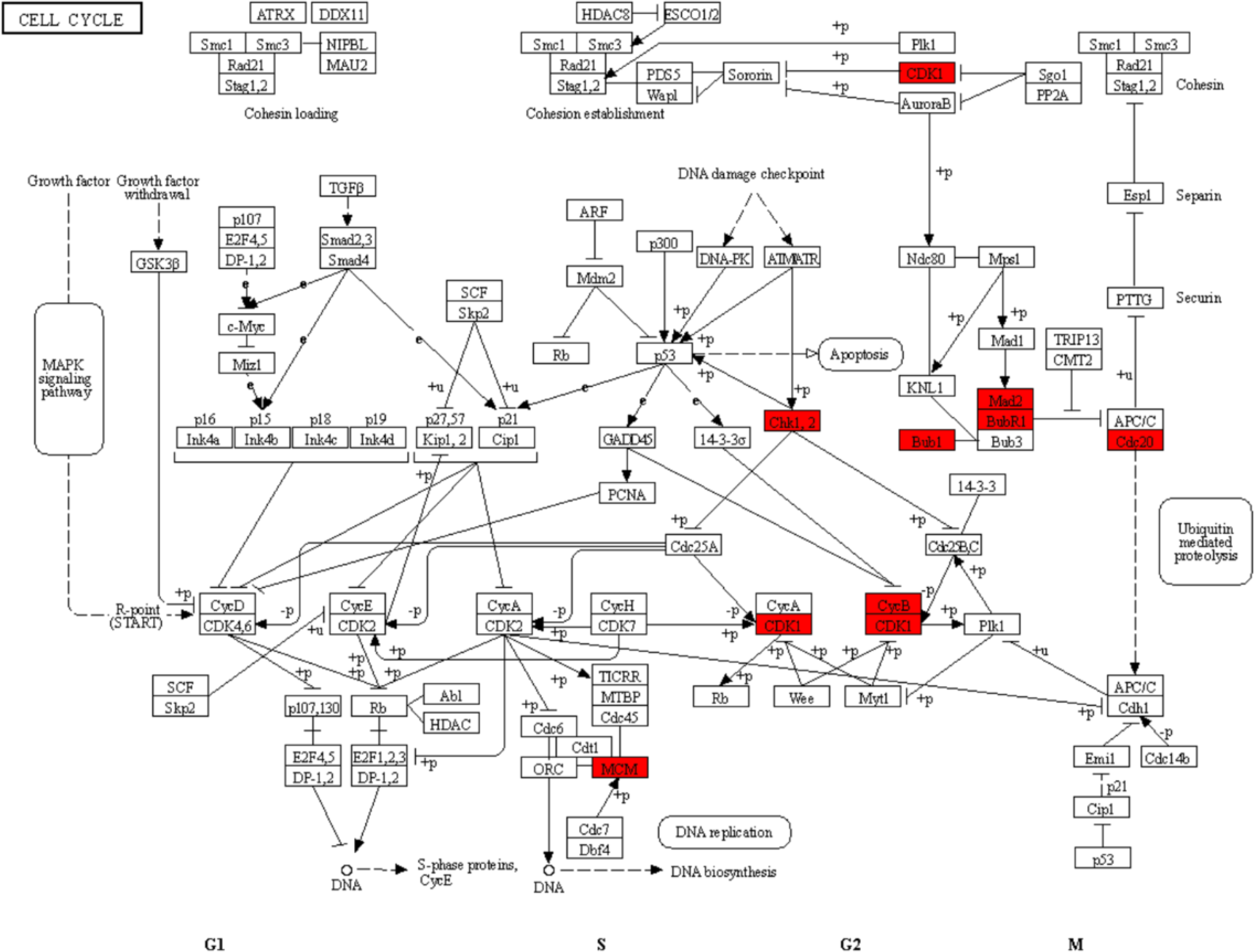
KEGG Pathway: Cell cycle (hsa04110) with annotations of downregulated DEGs. The cell cycle pathway includes four main phases: Gap 1 (G1), DNA replication synthesis (S), Gap 2 (G2), and Mitosis (M). Related pathways include apoptosis, MAPK signaling pathway, DNA replication, and Ubiquitin mediated proteolysis. The downregulated genes (highlighted red) include cyclin dependent kinase 1 (CDK1), checkpoint kinase 1 &2 (Chk1,2), mitotic checkpoint serine/threonine kinase (Bub1) and mitotic checkpoint serine/threonine kinase B (BubR1), mitotic arrest deficient 2 (Mad2), cell division cycle 20 (Cdc20), cyclin B3 (CycB), and minichromosome maintenance complex component 2 (MCM).

**Figure S9.**
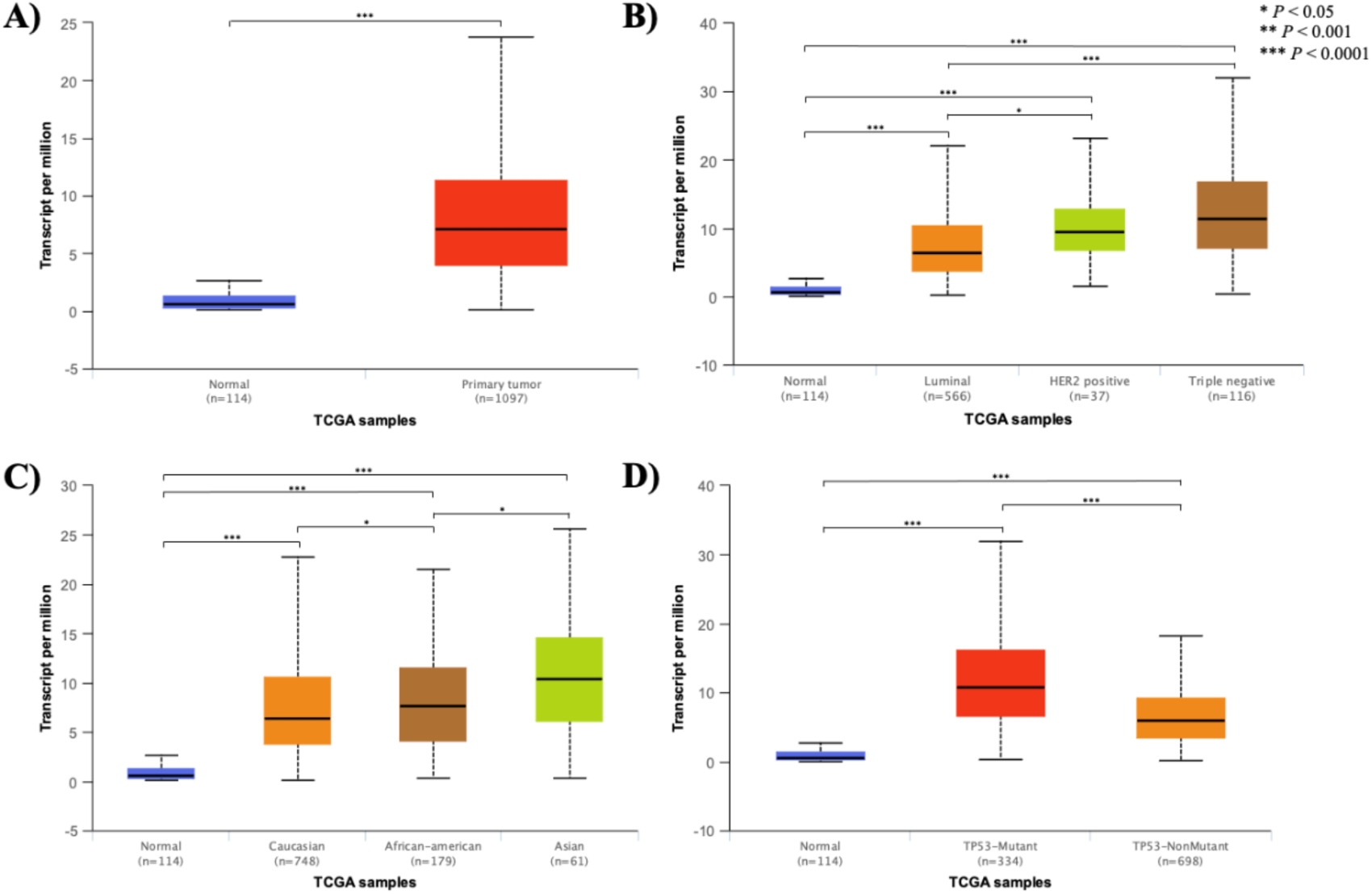
Comparative analysis of *HMMR* expression profile dynamics in breast cancer cell lines (performed using the UALCAN portal). The box plots and statistics were generated with expression data from The Cancer Genome Atlas (TCGA) database. All boxes in the diagram with asterisks signify significant differential expression, such that one asterisk denotes a *P* value < 0.05, two asterisks denote a *P* value < 0.01, and three asterisks denote a *P* value < 0.001. The box plots demonstrate varying expression levels in normal tissue (blue) and (A) breast invasive carcinoma tissue (red), (B) breast invasive carcinoma tissue of the luminal (orange), HER2 positive (green), and triple negative (brown) subtypes, (C) breast invasive carcinoma tissue in Caucasian (orange), African American (brown), and Asian (green) patients, and (D) breast invasive carcinoma tissue depending on TP53 mutation (red) status or TP53 non-mutant (orange) samples.

**Figure S10.**
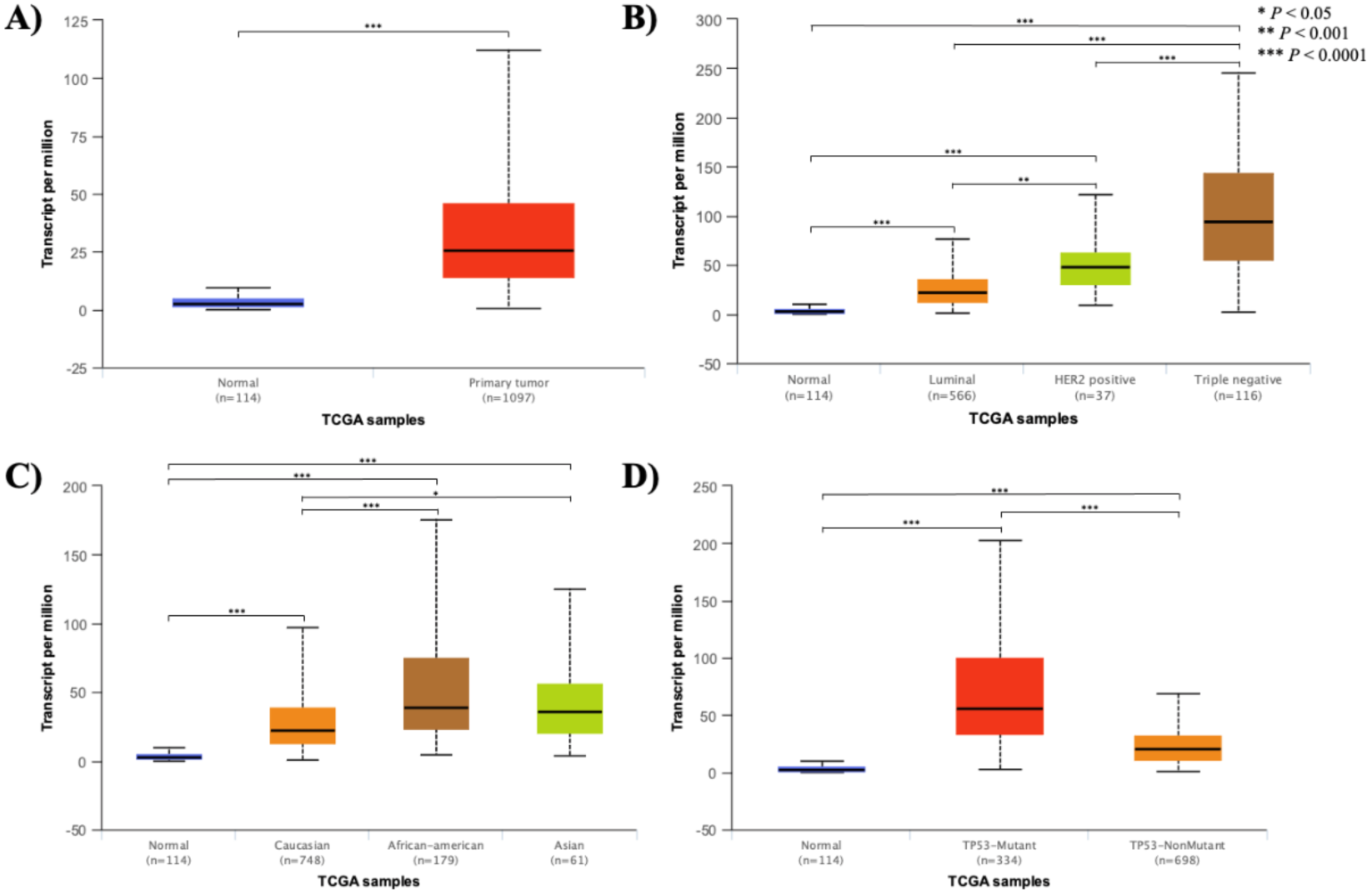
Comparative analysis of *CDC20* expression profile dynamics in breast cancer cell lines (performed using the UALCAN portal). The box plots and statistics were generated with expression data from The Cancer Genome Atlas (TCGA) database. All boxes in the diagram with asterisks signify significant differential expression, such that one asterisks denotes a *P* value < 0.05, two asterisks denote a *P* value < 0.01, and three asterisks denote a *P* value < 0.001. The box plots demonstrate varying expression levels in normal tissue (blue) and (A) breast invasive carcinoma tissue (red), (B) breast invasive carcinoma tissue of the luminal (orange), HER2 positive (green), and triple negative (brown) subtypes, (C) breast invasive carcinoma tissue in Caucasian (orange), African American (brown), and Asian (green) patients, and (D) breast invasive carcinoma tissue depending on TP53 mutation (red) status or TP53 non-mutant (orange) samples.

**Figure S11.**
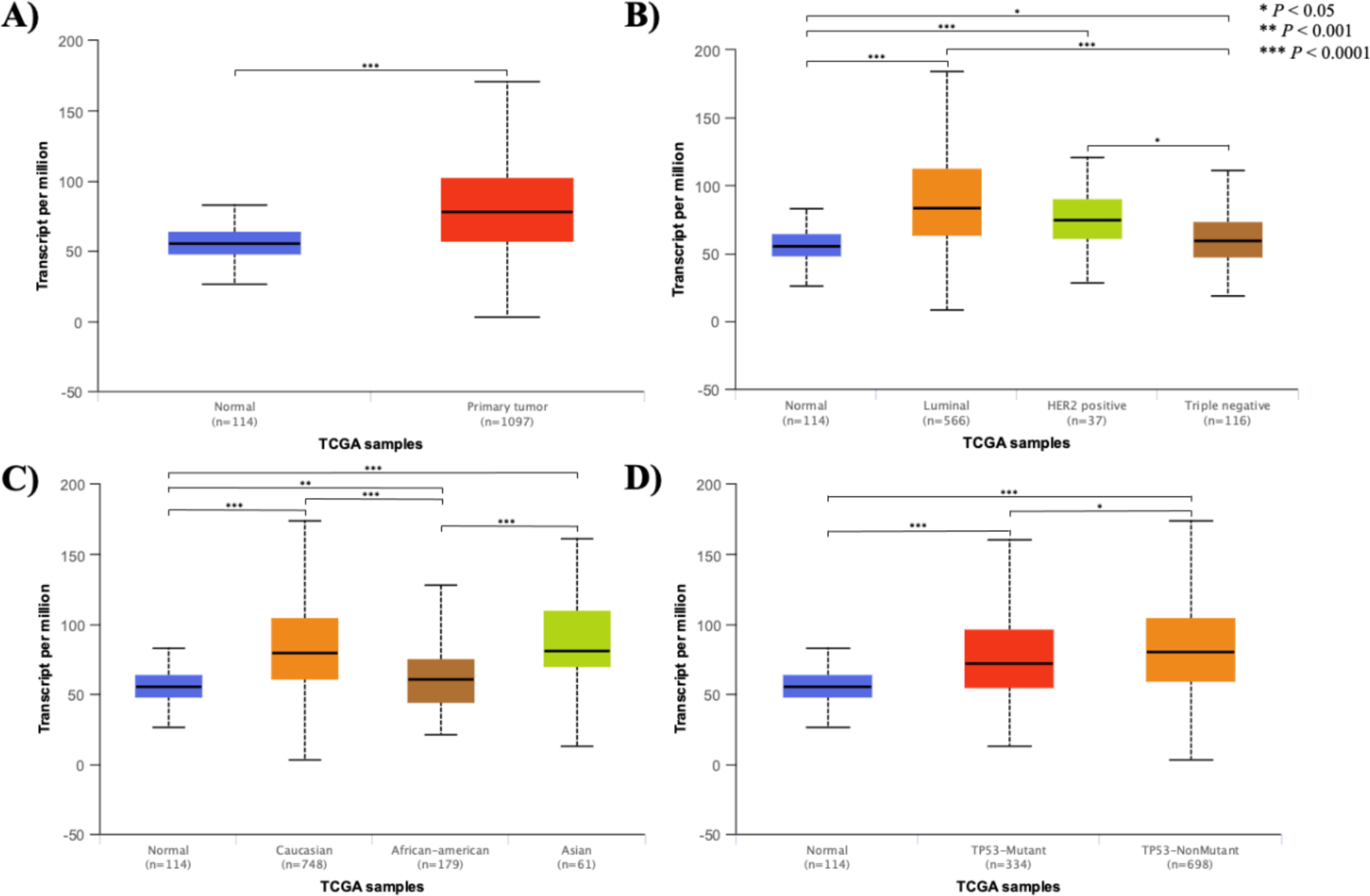
Comparative analysis of *HSPA4* expression profile dynamics in breast cancer cell lines (performed using the UALCAN portal). The box plots and statistics were generated with expression data from The Cancer Genome Atlas (TCGA) database. All boxes in the diagram with asterisks signify significant differential expression, such that one asterisks denotes a *P* value < 0.05, two asterisks denote a *P* value < 0.01, and three asterisks denote a *P* value < 0.001. The box plots demonstrate varying expression levels in normal tissue (blue) and (A) breast invasive carcinoma tissue (red), (B) breast invasive carcinoma tissue of the luminal (orange), HER2 positive (green), and triple negative (brown) subtypes, (C) breast invasive carcinoma tissue in Caucasian (orange), African American (brown), and Asian (green) patients, and (D) breast invasive carcinoma tissue depending on TP53 mutation (red) status or TP53 non-mutant (orange) samples.

**Figure S12.**
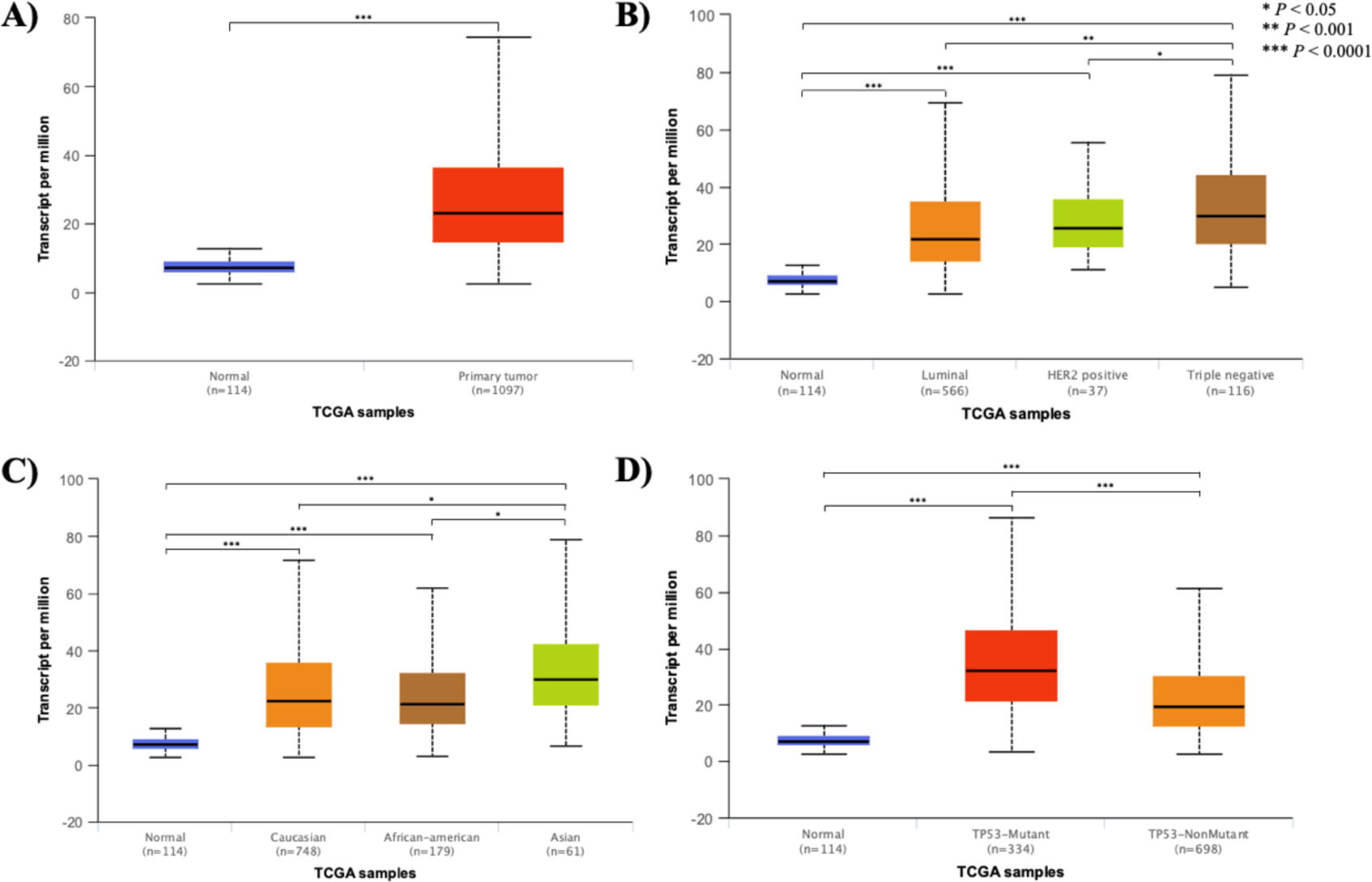
Comparative analysis of *RACGAP1* expression profile dynamics in breast cancer cell lines (performed using the UALCAN portal). The box plots and statistics were generated with expression data from The Cancer Genome Atlas (TCGA) database. All boxes in the diagram with asterisks signify significant differential expression, such that one asterisks denotes a *P* value < 0.05, two asterisks denote a *P* value < 0.01, and three asterisks denote a *P* value < 0.001. The box plots demonstrate varying expression levels in normal tissue (blue) and (A) breast invasive carcinoma tissue (red), (B) breast invasive carcinoma tissue of the luminal (orange), HER2 positive (green), and triple negative (brown) subtypes, (C) breast invasive carcinoma tissue in Caucasian (orange), African American (brown), and Asian (green) patients, and (D) breast invasive carcinoma tissue depending on TP53 mutation (red) status or TP53 non-mutant (orange) samples.

**Figure S13.**
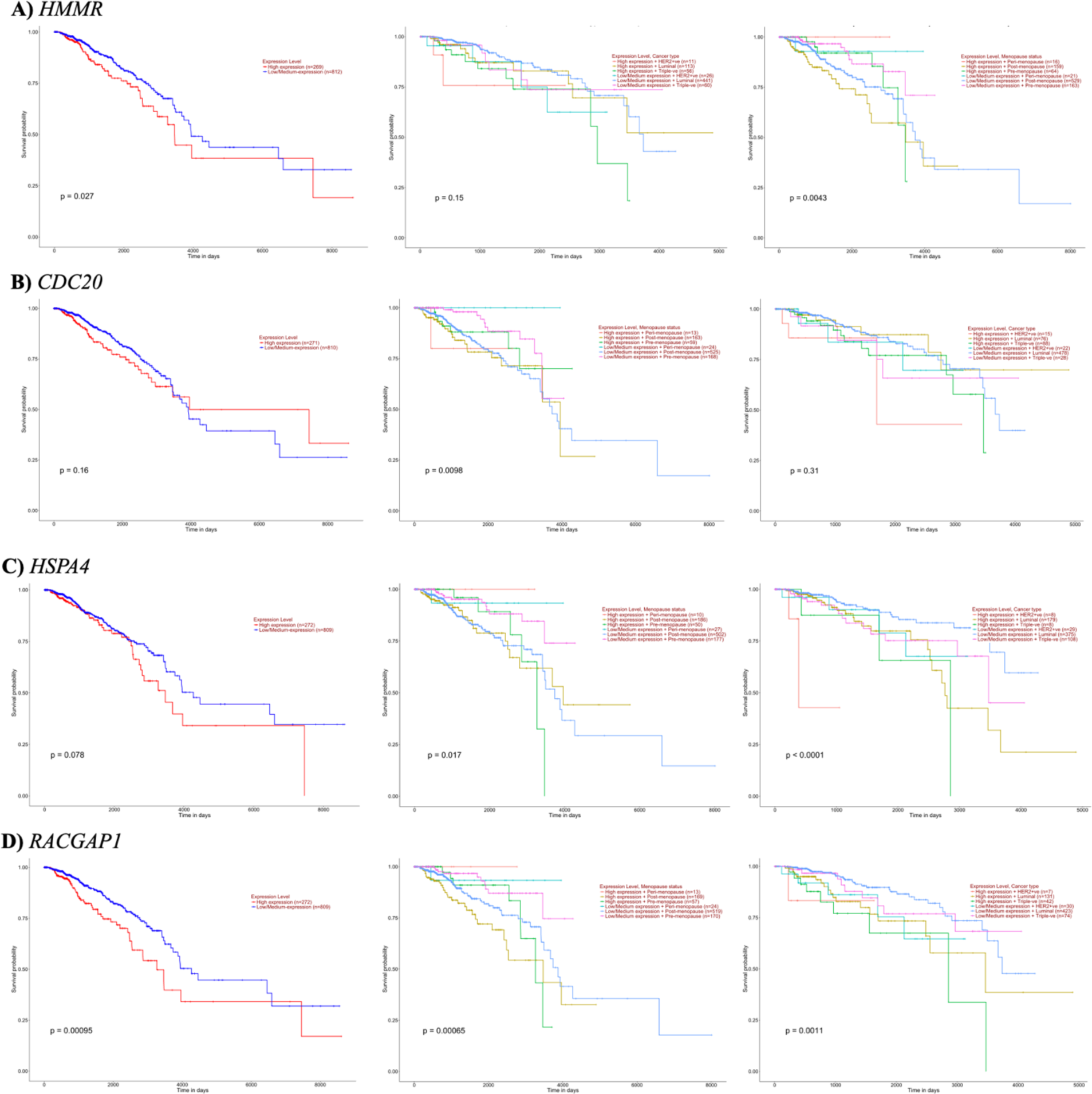
Survival analysis to determine prognostic effect in breast invasive carcinoma (performed using the UALCAN portal). The y-axis and x-axis show the survival probability (decimal) over time (in days), respectively. The survival plots show the correlation of high or low gene expression of (A) *HMMR*, (B) *CDC20*, (C) *HSPA4*, and (D) *RACGAP1* with survival rates in breast cancer overall, based on patient race, and based on menopause status.

**Figure S14.**
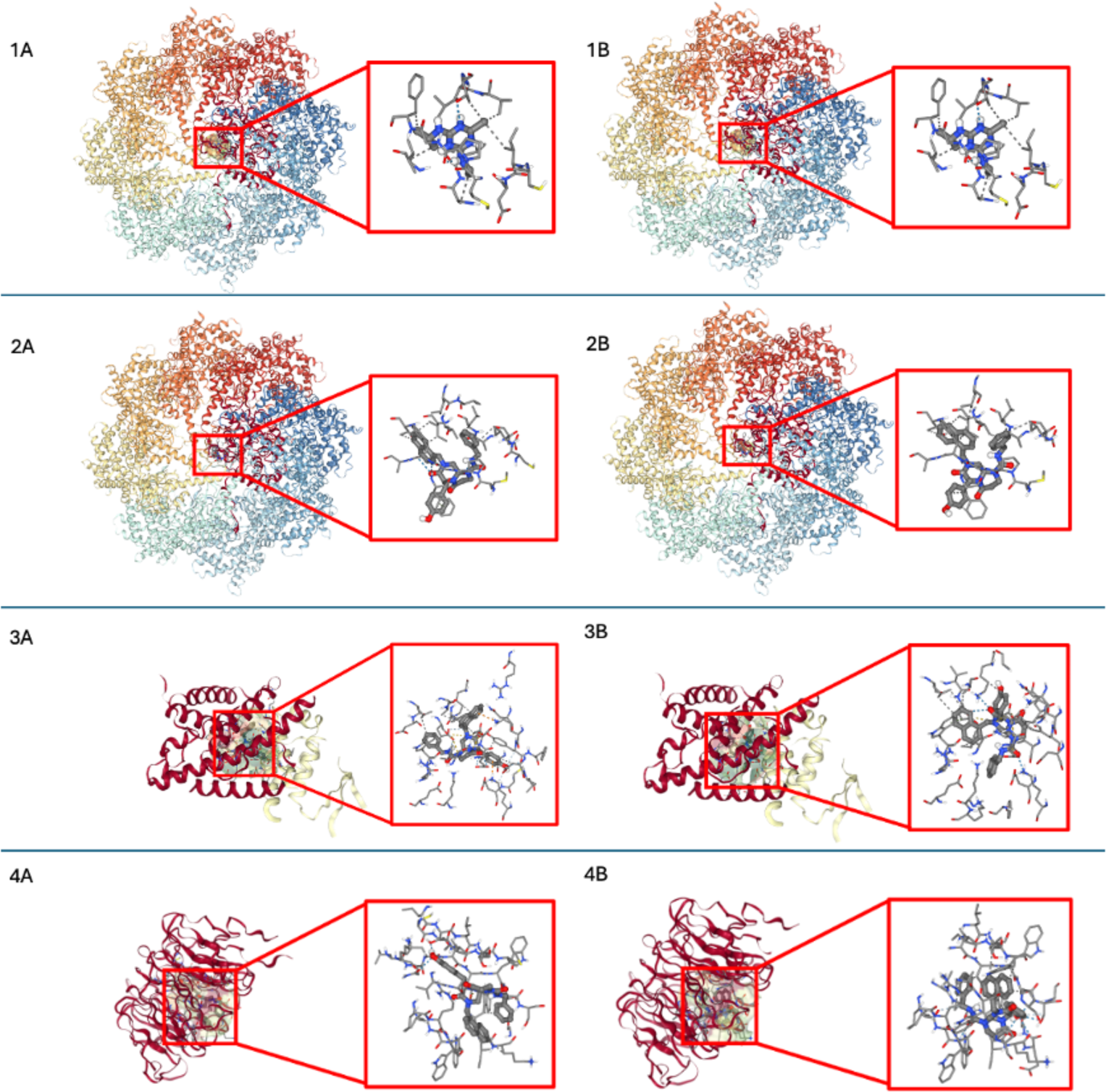
Seamdock results showing the (1A) molecular docking position of Altrazine (PubChem CID 2256) in 7TTT for Pose 1 with binding affinity of -5.3 kcal/mol, (1B) molecular docking position of Altrazine (PubChem CID 2256) in 7TTT (*CCT3* protein) for Pose 2 with binding affinity of -5.6 kcal/mol, (2A) molecular docking position of ICG 001 (PubChem CID 11238147) in 7TTT (*CCT3* protein) for Pose 1 with binding affinity of -9.3 kcal/mol, and (2B) molecular docking position of ICG 001 (PubChem CID 11238147) in 7TTT (*CCT3* protein) for Pose 2 with binding affinity of -8.9 kcal/mol. (3A) molecular docking position of ICG001 (PubChem CID 11238147) docked to 4IYP (*PPP2CA* protein) in Pose 1 with binding affinity of -10.2 kcal/mol. (3B) molecular docking position of ICG001 (PubChem CID 11238147) docked to 4IYP (*PPP2CA* protein) in Pose 2 with binding affinity of -9.3 kcal/mol. (4A) molecular docking position of ICG001 (PubChem CID 11238147) docked to 4GGC (*CDC20* protein) in Pose 1 with binding affinity of -10.9 kcal/mol (4B) molecular docking position of ICG001 (PubChem CID 11238147) docked to 4GGC (*CDC20* protein) in Pose 2 with binding affinity of -10.4 kcal/mol.

### Supplementary Tables

**Table S1.** Gene Ontology (GO) enrichment analysis for all DEGs. . For each Term ID, the ontology, description, FE, count, p-value (pval), and adjusted p-value (padj) are given.

| Ontology | Term ID | Description | FE | Count | pval | padj (FDR) |
| --- | --- | --- | --- | --- | --- | --- |
| BP | GO:0006270 | DNA replication initiation | 6.88 | 13 | 9.35E-08 | 2.58E-05 |
| BP | GO:0000070 | mitotic sister chromatid segregation | 6.80 | 17 | 5.61E-10 | 3.26E-07 |
| BP | GO:0007059 | chromosome segregation | 4.88 | 28 | 4.58E-12 | 5.46E-09 |
| BP | GO:0007052 | mitotic spindle organization | 4.85 | 19 | 2.52E-08 | 9.58E-06 |
| BP | GO:0006913 | nucleocytoplasmic transport | 4.75 | 17 | 2.31E-07 | 5.90E-05 |
| BP | GO:0090307 | mitotic spindle assembly | 4.70 | 13 | 1.07E-05 | 1.91E-03 |
| BP | GO:0007080 | mitotic metaphase plate congression | 4.58 | 13 | 1.40E-05 | 2.39E-03 |
| BP | GO:0042274 | ribosomal small subunit biogenesis | 4.26 | 21 | 4.53E-08 | 1.47E-05 |
| BP | GO:0032508 | DNA duplex unwinding | 3.95 | 16 | 7.49E-06 | 1.41E-03 |
| BP | GO:0006364 | rRNA processing | 3.70 | 34 | 7.46E-11 | 6.68E-08 |
| BP | GO:0051301 | cell division | 3.65 | 95 | 7.65E-29 | 2.74E-25 |
| BP | GO:0006260 | DNA replication | 3.43 | 28 | 2.68E-08 | 9.58E-06 |
| BP | GO:0000278 | mitotic cell cycle | 3.33 | 34 | 1.38E-09 | 6.19E-07 |
| BP | GO:0032543 | mitochondrial translation | 3.24 | 21 | 5.13E-06 | 1.08E-03 |
| BP | GO:0000398 | mRNA splicing, via spliceosome | 3.10 | 41 | 2.32E-10 | 1.66E-07 |
| BP | GO:0006281 | DNA repair | 2.94 | 59 | 1.55E-13 | 2.78E-10 |
| BP | GO:0008380 | RNA splicing | 2.69 | 38 | 5.90E-08 | 1.76E-05 |
| BP | GO:0007049 | cell cycle | 2.41 | 59 | 6.37E-10 | 3.26E-07 |
| BP | GO:0006325 | chromatin organization | 2.24 | 43 | 1.32E-06 | 2.95E-04 |
| BP | GO:0006974 | cellular response to DNA damage stimulus | 2.23 | 44 | 1.11E-06 | 2.66E-04 |
| BP | GO:0006338 | chromatin remodeling | 2.05 | 46 | 6.13E-06 | 1.22E-03 |
| CC | GO:0000796 | condensin complex | 11.53 | 6 | 5.47E-05 | 1.47E-03 |
| CC | GO:0000794 | condensed nuclear chromosome | 6.30 | 16 | 8.27E-09 | 4.60E-07 |
| CC | GO:0071007 | U2-type catalytic step 2 spliceosome | 5.63 | 11 | 1.18E-05 | 3.67E-04 |
| CC | GO:0000775 | chromosome, centromeric region | 4.89 | 21 | 3.61E-09 | 2.34E-07 |
| CC | GO:0032040 | small-subunit processome | 4.78 | 23 | 9.02E-10 | 6.38E-08 |
| CC | GO:0071005 | U2-type precatalytic spliceosome | 4.30 | 14 | 1.29E-05 | 3.86E-04 |
| CC | GO:0000776 | kinetochore | 4.20 | 44 | 6.93E-16 | 1.35E-13 |
| CC | GO:0005643 | nuclear pore | 4.09 | 25 | 4.72E-09 | 2.82E-07 |
| CC | GO:0005640 | nuclear outer membrane | 3.97 | 16 | 7.41E-06 | 2.51E-04 |
| CC | GO:0005637 | nuclear inner membrane | 3.76 | 22 | 2.27E-07 | 9.31E-06 |
| CC | GO:0071013 | catalytic step 2 spliceosome | 3.59 | 21 | 1.01E-06 | 3.74E-05 |
| CC | GO:0015030 | Cajal body | 3.55 | 18 | 8.49E-06 | 2.75E-04 |
| CC | GO:0005681 | spliceosomal complex | 3.35 | 29 | 2.60E-08 | 1.26E-06 |
| CC | GO:0000781 | chromosome, telomeric region | 3.25 | 37 | 5.43E-10 | 4.22E-08 |
| CC | GO:0005694 | chromosome | 3.24 | 54 | 3.58E-14 | 5.58E-12 |
| CC | GO:0005635 | nuclear envelope | 3.18 | 42 | 6.38E-11 | 5.51E-09 |
| CC | GO:0031965 | nuclear membrane | 2.62 | 44 | 1.12E-08 | 5.81E-07 |
| CC | GO:0072686 | mitotic spindle | 2.58 | 24 | 4.93E-05 | 1.37E-03 |
| CC | GO:0005813 | centrosome | 2.28 | 84 | 2.50E-12 | 2.77E-10 |
| CC | GO:0005874 | microtubule | 2.18 | 46 | 1.30E-06 | 4.61E-05 |
| CC | GO:0005654 | nucleoplasm | 2.05 | 538 | 1.87E-72 | 1.45E-69 |
| CC | GO:0005743 | mitochondrial inner membrane | 2.04 | 64 | 9.84E-08 | 4.25E-06 |
| CC | GO:0016607 | nuclear speck | 2.01 | 57 | 7.71E-07 | 3.00E-05 |
| MF | GO:0061608 | nuclear import signal receptor activity | 6.47 | 9 | 3.16E-05 | 3.11E-03 |
| MF | GO:0017056 | structural constituent of nuclear pore | 5.45 | 11 | 1.50E-05 | 1.81E-03 |
| MF | GO:0003678 | DNA helicase activity | 3.94 | 17 | 3.50E-06 | 5.42E-04 |
| MF | GO:0003777 | microtubule motor activity | 3.83 | 16 | 1.07E-05 | 1.45E-03 |
| MF | GO:0004386 | helicase activity | 3.76 | 22 | 1.95E-07 | 4.24E-05 |
| MF | GO:0003697 | single-stranded DNA binding | 2.80 | 22 | 3.01E-05 | 3.11E-03 |
| MF | GO:0016887 | ATPase activity | 2.74 | 80 | 2.82E-16 | 7.64E-14 |
| MF | GO:0042393 | histone binding | 2.63 | 35 | 3.50E-07 | 6.33E-05 |
| MF | GO:0003723 | RNA binding | 2.40 | 245 | 2.67E-40 | 2.89E-37 |

\*FE, Fold Enrichment; BP, biological process; CC, cellular component; MF, molecular function

**Table S2.**
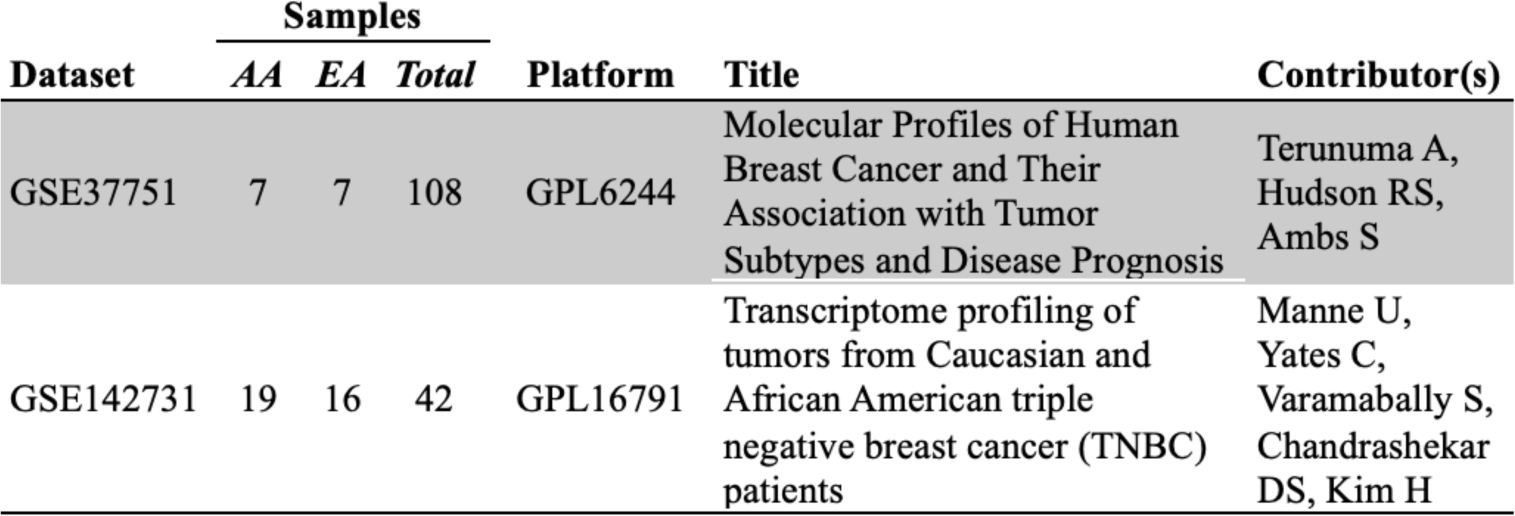
GEO datasets with sample numbers, title, and contributors.

